# Learning the Language of Phylogeny with MSA Transformer

**DOI:** 10.1101/2024.12.18.629037

**Authors:** Ruyi Chen, Gabriel Foley, Mikael Bodén

**Affiliations:** School of Chemistry and Molecular Biosciences, The University of Queensland, Brisbane, QLD 4067, Australia

## Abstract

Classical phylogenetics assumes site independence, potentially overlooking epistasis. Protein language models capture dependencies in conserved structural and functional domains across the protein universe. Here, we ask whether MSA Transformer, which takes a multiple sequence alignment (MSA) as input, captures evolutionary distance and to what extent its representations reflect epistasis in protein sequence evolution, neither of which are explicitly available during training. Systematic shuffling of natural and simulated MSAs demonstrates the model exploits column-wise conservation to distinguish phylogenetic relationships. Using internal embeddings, we reconstruct trees that are markedly consistent with trees generated by maximum likelihood inference. Applying this approach to both the RNA-dependent RNA polymerase of RNA viruses and the nucleo-cytoplasmic large DNA virus domain, we recover both established and novel evolutionary relationships. We conclude that MSA Transformer complements, rather than replaces, classical inference for more accurate histories of protein families.

## 1. Introduction

The growth in available biological sequences and their alignments, which reflect homology from common ancestry, enables the inference of phylogenetic relationships among diverse genomes or genes.^1^ The classical phylogenetics approach hitherto falls into several categories, including distance-based methods, such as Neighbor Joining (NJ),^2^ and character-based methods, including Maximum Likelihood (ML).^3^ While distance-based approaches progressively identify and merge pairs of taxonomic units for which distances are available or can be calculated,^4–6^ the ML approach employs evolutionary models to single out trees that assign the greatest likelihood to sequence characters when placed at their tips.^7,8^ As such, a phylogenetic tree depicts the evolutionary relationships between these sequences, hypothesising their shared ancestry. However, a general assumption of classical methods is that each position within an MSA evolves independently without influence from other positions.^1,9^ Epistasis manifests in a dependence between evolutionary outcomes and sequence context^10–12^ and upends such assumptions, potentially undermining accuracy of current phylogenetic inference methods.^13–15^

Protein Language Models (PLMs) *learn* dependencies *between* sites within structural and functional domains observed across diverse protein data,^16,17^ some via masked language modeling objectives.^18–22^ Typically, approximately 15% of amino acids are randomly masked, and model parameters are trained to recover those tokens given contextual information, potentially capturing complex dependencies between them.^19,23^ Similar to classical phylogenetics, several deep learning models,^24,25^ including MSA Transformer,^22^ AlphaFold’s EvoFormer,^26^ and EvoDiff-MSA,^27^ utilise MSAs of homologous proteins as inputs. The MSA Transformer is based on the BERT architecture,^28^ which incorporates interleaved row and column attention mechanisms across the sequences in the MSA; the model is trained on 26 million MSAs sourced from the UniRef50 database^29^ **(Figure 1A)**. The MSA Transformer has been applied to successfully predict residue contacts,^22^ protein structure,^30^ and the effect of mutations.^20^ Rao et al. showed that combinations of (tied) row attention heads of the MSA Transformer capture structural contacts in proteins.^22^ Lupo et al. noticed that combinations of MSA Transformer’s column attention heads correlate with the Hamming distance between input sequences, suggesting their application in tracing evolutionary lineages of proteins.^31^

**Figure 1:**
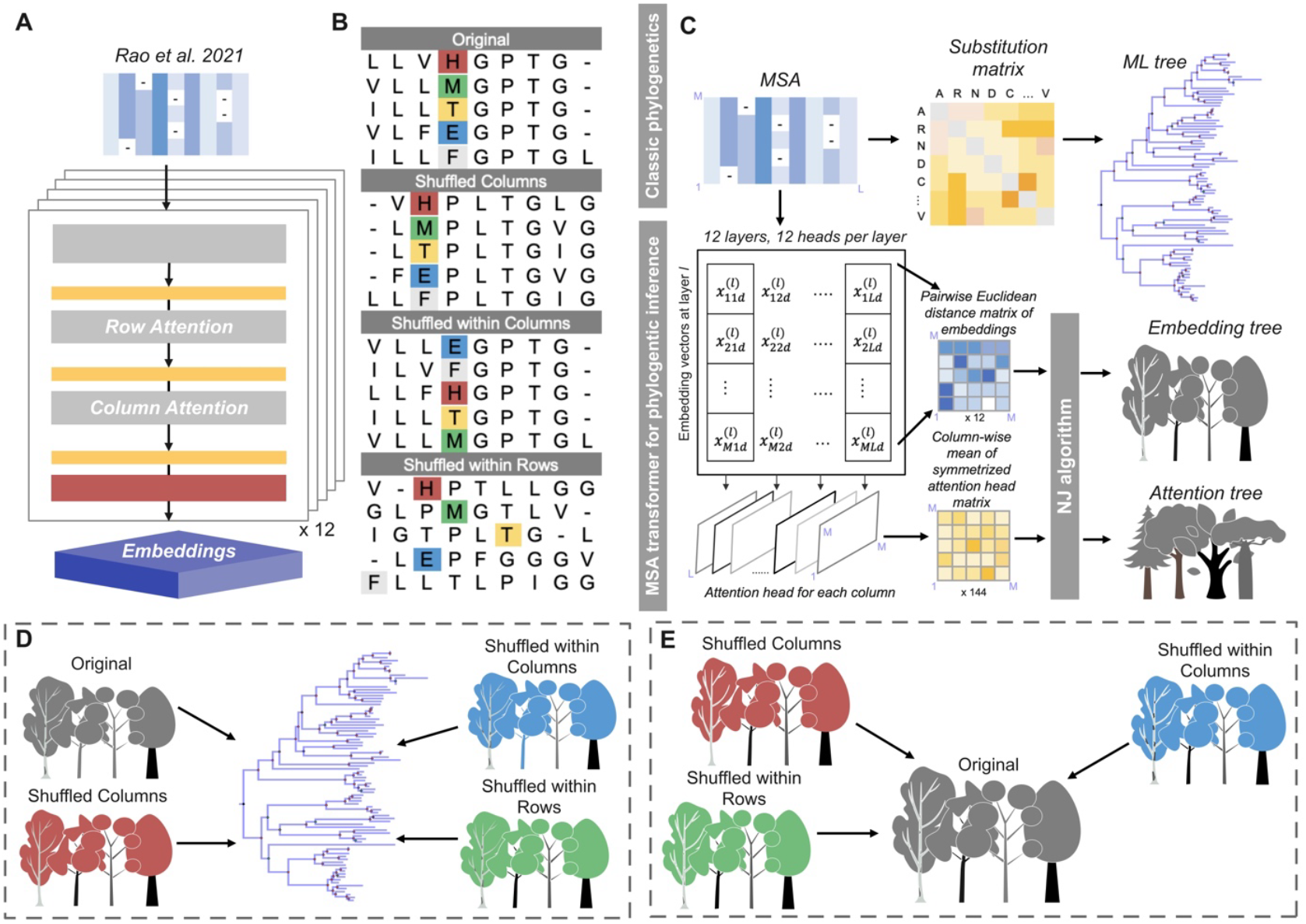
Overview of phylogenetic inference using MSA Transformer. **(A)** The structure of the MSA Transformer. **(B)** Four distinct MSAs are used in this study, with specific colors highlighting certain positions to illustrate changes in the alignment. **(C)** The general workflow for phylogenetic inference by MSA Transformer compared to likelihood-based methods. An MSA was sent into this model to extract embeddings and column attention across layers. Pairwise Euclidean distance matrices 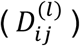 were computed for the three-dimensional embeddings at each layer. Additionally, 144 CMSCAH matrices 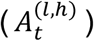 were generated as described in the “STAR Methods - Spearman rank correlation coefficients with evolutionary distances.” These matrices were then used as distance matrices in the NJ algorithm to reconstruct phylogenetic trees. **(D)** Exploration of the consistency between traditional phylogenetic trees and inferred trees constructed using four distinct MSAs. **(E)** Exploration of the factors in natural sequences distinguished by MSA Transformer. We evaluated the consistency of embedding trees constructed using the original MSA in comparison with those generated from three shuffled MSAs.

Here, we ask whether MSA Transformer captures evolutionary distance and to what extent its representations reflect epistasis in protein sequence evolution, even though neither of these features are explicitly available during training. We investigate the model’s reliance on information available in columns as opposed to rows in the MSA, by systematically shuffling its content (**Figure 1B**). We then use MSA Transformer on both natural and simulated MSAs (where epistasis is strictly absent) to reconstruct entire phylogenetic trees with implied ancestral branchpoints (**Figure 1C**) and assess their consistency with trees from maximum likelihood approaches (**Figure 1D**). To further probe which factors in natural sequences are distinguished by MSA Transformer, we compare the trees generated from shuffled alignments with those from the unshuffled alignment (**Figure 1E**). Finally, we demonstrate the complementarity of using MSA Transformer to classical phylogenetic inference by reconstructing phylogenetic trees for the RNA virus RNA-dependent RNA polymerase (RdRp) and DNA viruses from the Nucleo-Cytoplasmic Large DNA Virus (NCLDV) domain; we find both novel and previously known evolutionary relationships are available from a “non-classical” approach with different computational requirements.

## 2. Results

### 2.1 The MSA Transformer captures evolutionary distances

Lupo et al. used the Column-wise Means of the Symmetrised Column Attention Heads (CMSCAHs) from MSA Transformer, obtained by symmetrising and averaging attention values across all columns, as features for a logistic regression model to predict pairwise Hamming distances in the input MSA.^31^ This approach demonstrates how the column attention mechanism relates to phylogenetic information. Defined as the number of substitutions needed to change one sequence into another, the Hamming distance does not account for the continuous nature of evolution and the possibility of repeated mutations occurring at the same position.^32,33^ It is also not clear the extent to which embeddings and phylogeny correlate more broadly. To address these concerns, we (1) extracted evolutionary distances between pairs of sequences from two kinds of phylogenetic trees-NJ and ML-sampled across the protein universe,^34^ (2) calculated Euclidean distances (cosine also suitable)^35^ from embeddings of the MSA Transformer, along with CMSCAHs at different layers, and (3) benchmarked the concordance between (1) and (2) by calculating Spearman rank correlation coefficients (Spearman’s rho). We acquired seed alignments from 20 distinct protein families, together with their NJ trees, curated by the Pfam database. We also inferred ML trees based on the same seed alignments (“STAR Methods - Pfam data”; **Table S1**).

An MSA aligns sequences within a shared positional (column) coordinate system, unveiling where homology exists and identifying patterns of conservation.^36^ We hypothesize that the MSA Transformer leverages column-wise conservation within an input MSA to infer phylogenetic relationships, thereby enabling its potential application in phylogenetic reconstruction. To evaluate this, conservation information encoded within each column was systematically examined through controlled experiments. We considered four versions of an MSA for each protein family: (1) the original MSA, (2) the shuffled-columns MSA, (3) the shuffled-within-columns MSA, and (4) the shuffled-within-rows MSA (**Figures 1B** and **S1**).

For each original MSA (1), which is obtained directly from Pfam, we produced the three versions (2-4), each of which retains the original overall amino acid composition (versions 2-3 were introduced by Rao et al.^22^) The shuffled-columns MSA randomly reorders columns and thereby alters cross-positional dependencies; however, internally each column is unchanged, preserving zero-order conservation and composition when stratified by row (i.e. sequence index). The shuffled-within-columns MSA retains the column order of the original MSA but randomly mixes amino acids *within* each column; the basis for calling a column conserved is thus retained (viewed independently from other positions), but columns mix content from different rows (i.e. sequence indices). We introduced a shuffled-within-rows MSA where values within each row are permuted, retaining composition when stratified by row but disrupting it when stratified by column, all relative to the original MSA.

For each protein family, we input the four MSAs into the MSA Transformer, extracting embeddings and column attention for each layer. Using the attention of the column and the embeddings extracted for each version of an MSA, we computed the CMSCAH^31^ and the pairwise Euclidean distances of their embeddings. From these, across all pairs of sequences within a protein family, we determined the Spearman’s rho to the additive distances indicated by the phylogenetic trees produced by NJ and ML (see “STAR Methods - Spearman rank correlation coefficients with evolutionary distances” for details). In plots and tests below, we used the Spearman’s rho of pairwise evolutionary distances computed from the NJ and ML trees as a reference-indicating a baseline margin of difference that can be explained by switching one classical inference method out for another. For statistical comparisons, shuffling was performed five times with different random seeds, recorded with the average Spearman’s rho, along with its standard deviation. To facilitate a direct comparison with the results of Lupo et al., we applied the same method (supervised prediction of Hamming distances^31^) to CMSCAHs derived from the original MSA. We summarized and presented the overall trends for all 20 protein families based on evolutionary distances obtained from the NJ tree (**Figures 2A** and **2B**), with detailed results for each family provided in **Figures S2-S3**.

**Figure 2:**
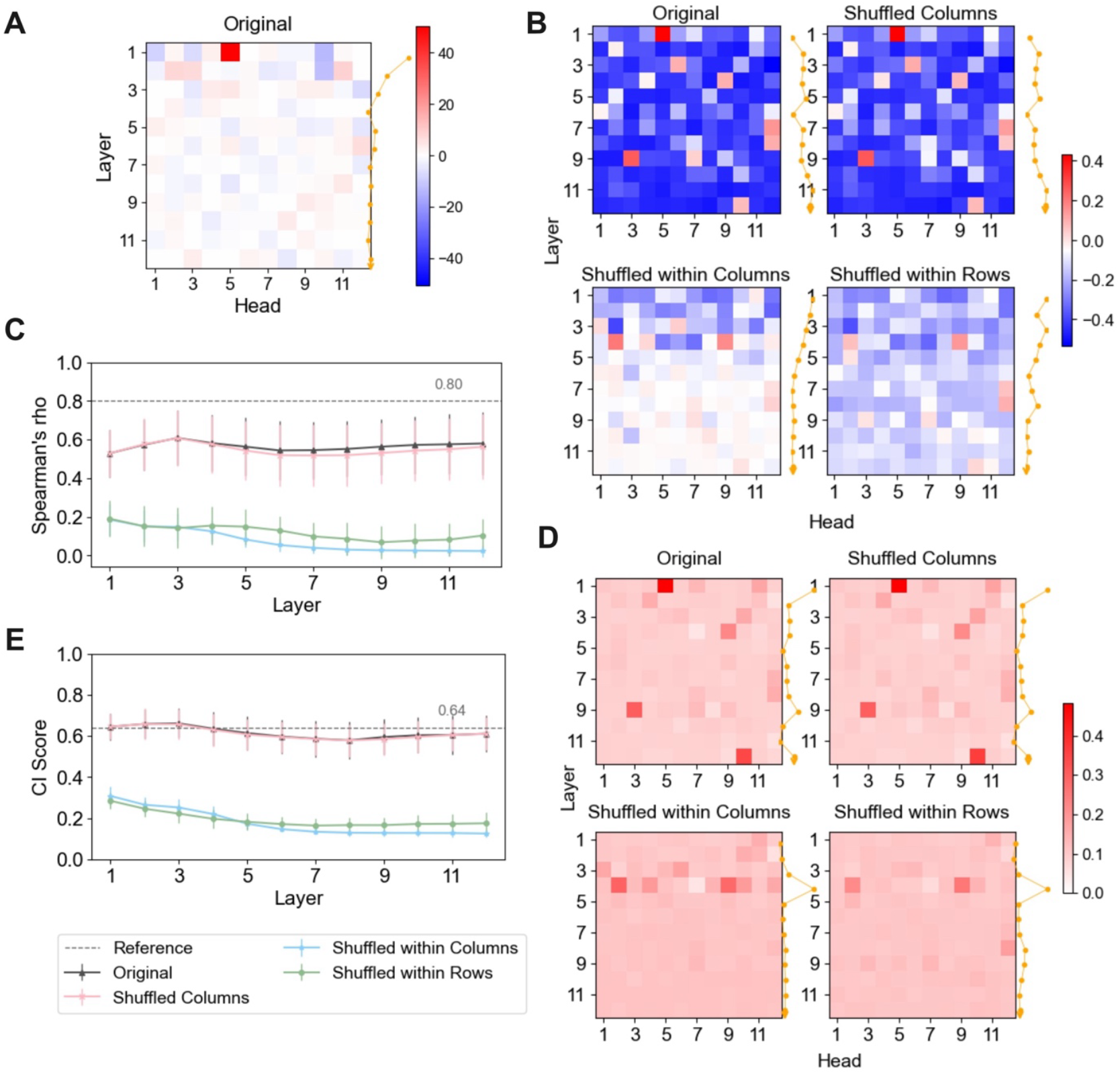
Phylogenetic signals captured by MSA Transformer in four MSAs. **(A)** Average regression coefficients for 144 CMSCAHs predicting pairwise Hamming distances, following the methods of Lupo et al.^31^ **(B)** Average Spearman’s rho between CMSCAHs and pairwise evolutionary distances from the NJ tree for four MSAs. **(C)** Average Spearman’s rho between pairwise Euclidean distances of embeddings and evolutionary distances from the NJ tree. (**D, E**) Average CI scores between (D) attention trees or (E) embedding trees and the NJ tree, respectively. The yellow line shows the average absolute value across heads for each layer, the error bars represent the standard deviation, and the gray dashed line marks the reference value between NJ and ML trees; all are averaged across 20 protein families.

#### CMSCAHs correlate with evolutionary distances

Lupo et al. demonstrated that the regression coefficients of a logistic model of CMSCAH combinations are consistently predictive of (Hamming) distances between sequences; moreover, they remained predictive for *different* protein families (**Figures 2A, S2A**, and **S3A**). Given that the number of differing positions indicates the rate of evolution over short time frames, Spearman’s rho between attention heads and the evolutionary distance unsurprisingly displayed the same trend across protein families (the first column in **Figures 2B, S2B-2C**, and **S3B-3C**). The fifth attention head in the first layer (i.e., Head 1-5) showed a strong positive Spearman’s rho with phylogenetic distance, while most column attention heads were *negatively* correlated. At the per-layer level, correlations with evolutionary distance became stronger (more negative) with increasing layer depth, which is opposite to the trend observed in **Figure 2A**.

#### Distances between pairs of sequences in the embedding space correlate positively with their phylogenetic distances

Compared to Head 1-5, the Euclidean distances of layer-specific embeddings between sequences in the original MSA had a greater positive Spearman’s rho with the distances extracted from the trees inferred by both NJ and ML (**Figures 2C** and **S4**). Distances in early-layer embeddings generally indicated stronger positive correlations with evolutionary distances compared to later layers but remained lower overall than correlations between the distances inferred by ML and NJ.

#### The capture of evolutionary distances mainly relies on conservation information within columns

The original MSA exposes the pattern of conservation in its entirety (including potential dependencies across columns); the shuffled-columns MSA maintains column conservation but disrupts dependencies across columns that may have been learned, the shuffled-within-columns maintains the skew in *composition* of amino acids within columns, and the shuffled-within-rows removes column conservation altogether. We observed that the shuffled-columns MSA produced Spearman’s rho values comparable to those of the original MSA when correlated with evolutionary distances (**Figures 2B-2C** and **S2**-**S4**). In contrast, the results from both shuffled-within-columns and shuffled-within-rows MSAs departed significantly from the first two but were comparable to each other. Given that the shuffled-within-columns MSA maintains zero-order conservation, while scrambling rows, we suggest that presenting conservation within columns is *necessary* but *not sufficient* to recover evolutionary distances in the MSA Transformer. Classical phylogenetic inference assumes independence between sites and would recover the *same* evolutionary distances for the original and the shuffled-columns MSA, implying that (like ML or NJ inference) MSA Transformer does *not* rely exclusively on similarities between the input and specific sequences it has seen *during training*.

### 2.2 MSA Transformer trees are similar to trees inferred by classical phylogenetics

Neighbor joining takes a distance matrix typically calculated from a sequence alignment; the distance matrix is amended for “internal” nodes, with distances based on the principle of additivity as pairs of less ancient nodes are progressively “collapsed”. Here, we used pairwise Euclidean distances determined by inspecting the MSA Transformer embedding space and attention heads as input to NJ, yielding “embedding trees” and “attention trees”, respectively (**Figure 1C**). To evaluate the ability of MSA Transformer trees to represent phylogenetic relationships, we employed Robinson Foulds (RF^37^) and Clustering Information (CI^38^) scores to measure topological tree similarity (“STAR Methods - Phylogenetic tree reconstruction and similarity comparison”). The similarity score between the NJ and ML trees was also calculated for each protein family, serving as a reference for comparison (family specific results are shown in **Figures S5-S9**). To capture trends, we aggregated results across 20 protein families based on the inferred tree and the NJ tree evaluated by CI score.

#### When the column attention head is positively correlated with the evolutionary distance, the corresponding attention tree is similar to the NJ and ML trees

We observed that attention heads with a negative correlation to evolutionary distances (**Figure 2B** and **S2-S3**) generally had low similarity with two phylogenetic trees (**Figures 2D** and **S5-S9**). The similarity scores of 144 CMSCAHs exhibited consistency across 20 protein families (**Figure S10**), as demonstrated by a high Pearson correlation (**Figure S11**).

#### Select attention trees are similar to NJ and ML trees

Head 1-5 was identified for its positive correlation with evolutionary distances (**Figure 2B**). The tree derived from Head 1-5 accurately reconstructed both NJ and ML trees (**Figures 2D** and **S5-S9**). Besides, Head 12-10, previously overlooked due to its weak concordance with distance metrics, also produced an attention tree that consistently recovered these two phylogenetic trees.

#### Embedding trees from early as opposed to later layers in the MSA Transformer are more similar to NJ and ML trees

We noted that a high Spearman’s rho for distances in embedding space does not necessarily imply improved tree similarity (**Figures S4** and **S12-S13**). In 60% of protein families, the maximum CI score (observed at early layers) between the embedding tree and the NJ tree either matched or exceeded the similarity between the NJ and ML tree, with the NJ v. ML similarity serving as our reference baseline (**Figure S12**). Across all 20 protein families analysed, embedding trees from the early layers exceeded the average baseline similarity (**Figure 2E**). It suggested that early-layer embeddings capture enough phylogenetic signal to reconstruct protein family trees with accuracy comparable to traditional methods. This observation aligns with previous research that final-layer representations are not always optimal, and that finetuning allows useful information from middle layers to be utilized by downstream tasks.^39^

#### To reproduce the topology of NJ and ML trees, MSA Transformer mainly relies on conservation information within columns

The MSA Transformer trees produced from the shuffled-within-column MSAs, where conservation is not indexed by sequence, and the shuffled-within-rows MSAs, which disrupt column indexing and essentially remove conservation, showed low similarity across all configurations (**Figures 2D** and **2E**). In contrast, shuffled-columns MSAs preserve column-wise conservation, facilitating the reconstruction of NJ and ML trees that are highly similar to those obtained from the original MSA.

### 2.3 The MSA Transformer primarily relies on column-wise conservation to infer phylogeny

We further investigate whether the MSA Transformer relies on column-wise conservation information to infer tree topology, focusing on embedding trees (**Figure 1E**). For all protein families, the similarity between embedding trees from the original MSA and shuffled-column MSA successively decreases across layers (much approaching 1 at the first layer) (**Figures 3A** and **S14**). This contrasts with comparisons of the layer-specific embedding trees (built from MSA versions 1-2) to the NJ tree, where similarity remained near-constant across layers (**Figures 2E** and **S12**). For instance, in Pfam PF14317, early-layer embedding trees were minimally affected by column reordering, while later-layer embeddings exhibited distinct topological changes (**Figure S15**). Furthermore, embedding trees derived from MSA versions 3-4 showed significantly lower similarity to those from the original MSA compared to trees generated from shuffled-column MSAs across all layers. These differences underscore that MSA Transformer primarily relies on column-wise conservation to infer tree topology and exhibits increased sensitivity to the biological veracity of sequences in later layers.

**Figure 3:**
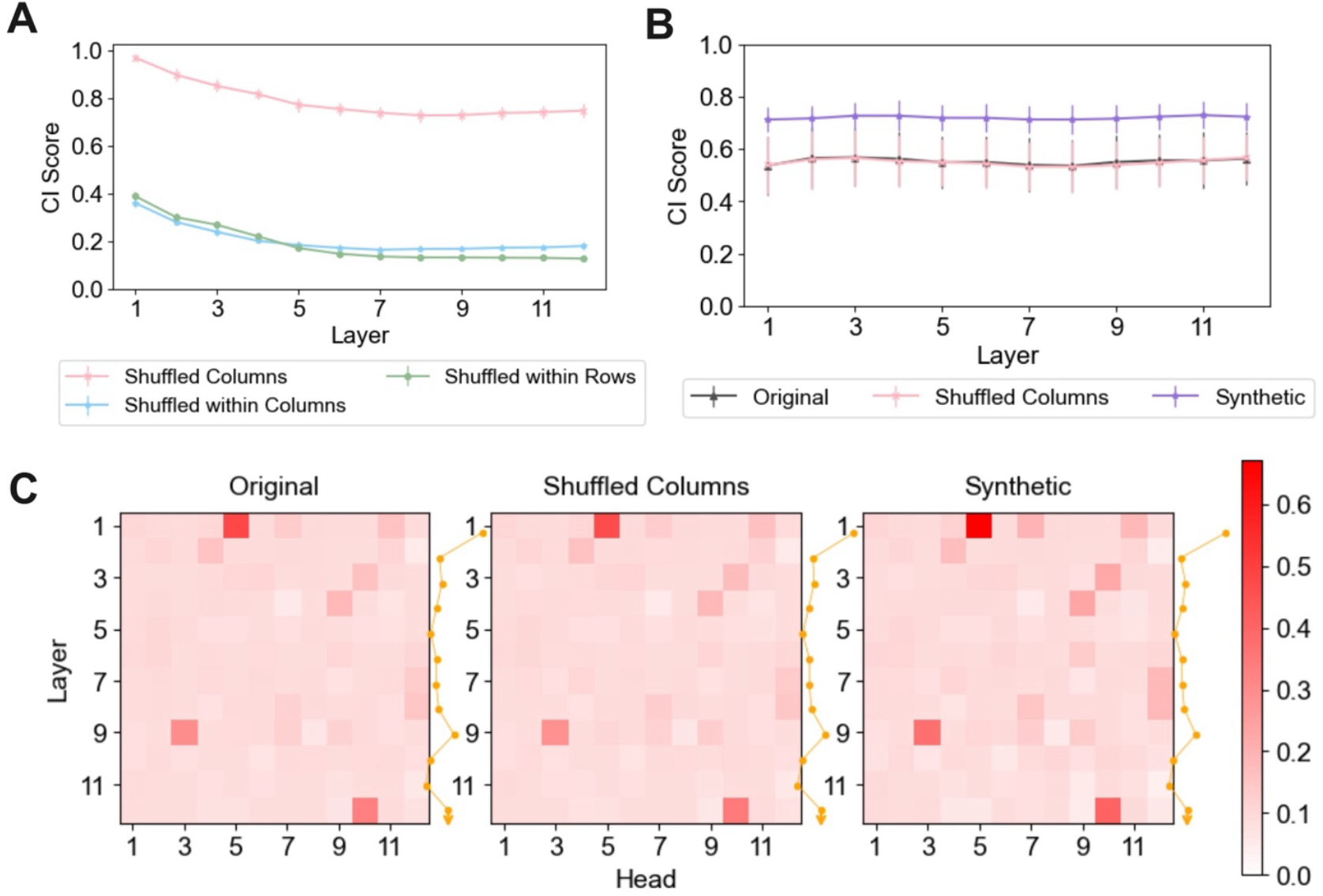
The MSA Transformer primarily relies on column-wise conservation information to infer phylogeny. (**A**) Layer-by-layer CI scores were compared between embedding trees from three types of shuffled MSAs and those from the original MSAs, averaged across 20 protein families. (**B, C**) CI scores of (B) embedding trees and (C) attention trees constructed from original, shuffled-columns, and synthetic MSAs were compared with those of their corresponding ML trees averaged across 20 protein families. For Pfam MSAs, only ML trees derived from the original MSA were used for comparison with the embedding trees. The yellow line shows the mean absolute CI scores across heads for each layer, and error bars indicate the standard deviation; both are averaged across 20 protein families.

Despite these observed similarities, MSA Transformer and classical methods differ fundamentally in their underlying mechanisms. Unlike MSA Transformer, classical phylogenetic inference does *not* rely on training; MSA Transformer is pre-trained with existing protein sequences, which *can* reflect epistasis (see section 2.4). It is thus important to acknowledge that deviations from NJ or ML trees do *not* imply that one is more (or less) accurate than the other. Moreover, assuming epistasis is present in the input MSA, no or minor differences between classical phylogenetic inference and an approach based on MSA Transformer would indicate that effects were *not* captured by either. Below, we use MSA Transformer on both natural (where epistasis *may* be present) and synthetic MSAs (where it is strictly absent) to address what complementarity can be discerned between the two classes of approaches.

We generated a controlled synthetic MSA for each protein family using TrAVIS, which is part of the GRASP-suite (see “STAR Methods - Synthetic MSAs” and **Tables S3-S4**). For each synthetic MSA, we applied the same methodology described before to construct ML and MSA Transformer trees, and compute similarity scores between the trees. (The differences among all MSA types used in our study are summarised in **Table S2**.) Here, we compared MSA Transformer’s ability to construct three MSA types that differ in biological meaning but retain column-wise conservation information.

#### The MSA Transformer distinguishes some aspects of biological meaning

Analysis of embedding trees from different layers revealed no significant differences between trees generated using MSA versions 1-2, compared to the ML trees (inferred by MSA version 1), for 55% protein families (Welch’s two-tailed t-test, *p* > 0.05). However, embedding trees based on synthetic MSAs were significantly more similar to their corresponding ML trees (inferred by synthetic MSAs) compared to embedding trees derived from Pfam MSAs and their ML trees (Welch’s one-tailed t-test, *p* < 0.001) (**Figures 3B** and **S16**). A notable exception included the CI score for a synthetic SPF07498, which was not markedly higher than that of the original PF07498 (**Figure S16B**). Trees constructed from most attention heads for synthetic data were similar to those for Pfam data when compared to their ML trees, except for the attention trees built from Heads 1-5 and 12-10 (**Figures 3C** and **S17-S20**). The synthetic attention tree for Head 1-5 improved similarity to its ML tree, with average RF and CI scores increasing by 136% and 48% across 20 protein families. In contrast, the Head 12-10 trees improved for only 75% of families.

With simulated data, the MSA Transformer can reconstruct phylogenetic trees that closely match those inferred by maximum likelihood methods, even though the data is not part of its training set and lacks epistasis. This indicates the model captures phylogenetic features beyond memorizing training examples, generalizing across protein families (see Tule et al.^40^ for complementary analyses). However, with real MSAs, the greater differences between MSA Transformer and classical inference suggest that these approaches yield qualitatively different results on complex biological data. Thus, embedding and attention trees may interpret evolutionary relationships in a more context-dependent way, though we cannot say which is more accurate.

### 2.4 The MSA Transformer recovers epistatic interactions

To test the MSA Transformer’s principal capability to detect epistatic effects and make them available for further processing, we collected the protein fitness deep mutational scanning (DMS) dataset, along with the wild-type MSA for the avGFP protein, described in Chen et al.^41^ We randomly selected 1,000 double mutants from that dataset. For each variant, we calculated the epistasis value and classified the variants as either epistatic or non-epistatic (“STAR Methods - Epistasis calculation and identification of epistatic interactions”). We extracted embeddings for the reference sequence at the mutated positions from both the wild-type and mutant MSAs. Using the 1,000 double mutant samples, we trained a ridge model on concatenated embeddings from the wild-type and mutant MSAs to predict epistasis from DMS dataset.

We observed a Pearson correlation of 0.0859 (*p* = 0.0066) between the predicted and experimentally measured epistasis. Despite the low correlation, the significant *p*-value implies that the MSA Transformer embeddings contain information pertinent to epistatic effects. Based on the predictions from the ridge model, we classified those double mutants as either epistatic or non-epistatic using the same criteria. The results indicate that the model can distinguish between non-epistatic and epistatic variants to a limited but statistically significant extent, and is particularly accurate in identifying non-epistatic mutations (**Figure S21**).

### 2.5 Accurate reconstruction of RNA virus RdRp domain

What evolutionary insights does the MSA Transformer embedding tree provide, and how consistent are they relative to those based on the ML tree? We next evaluate if embedding trees are able to recover phylogenies of species with better-known evolutionary histories, such as viruses.

Shi et al. previously conducted an extensive analysis of transcriptomes from over 220 invertebrate species across nine animal phyla, leading to the discovery of 1,445 RNA viruses.^42^ To position these newly discovered viruses within the broader context of known viral biodiversity, they constructed phylogenies using the RdRp domain, the only domain present in all RNA viruses. They reclassified viral families, orders, and genera into 16 distinct RNA virus clades, like “Orthomyxo” for Orthomyxoviridae and “Reo” for Reoviridae.

We acquired four well-annotated MSAs with their ML trees: the “Astro”, “Orthomyxo”, “Nido”, and “Reo” clades (the term “clade” is used here, but it should be noted that each represents an independent MSA and a tree, as detailed in “STAR Methods - RdRp and NCLDV data” and **Table S5**). Embedding trees derived from different layers exhibited varying degrees of similarity to traditional phylogenetic trees, necessitating further analysis to identify the particular layer for comparison. Similarity scores between embedding trees and ML trees were computed across layers for each clade (**Table S6**), revealing that layer 2 or 3 consistently achieved the highest RF scores and was thus selected. To facilitate the comparison of evolutionary relationships by topology, branch lengths were hidden. Here, we present the comparison of embedding trees with ML trees for the “Reo” and “Nido” clades (**Figure 4**), which feature larger ML trees. Comprehensive results are provided in **Figure S22**.

**Figure 4:**
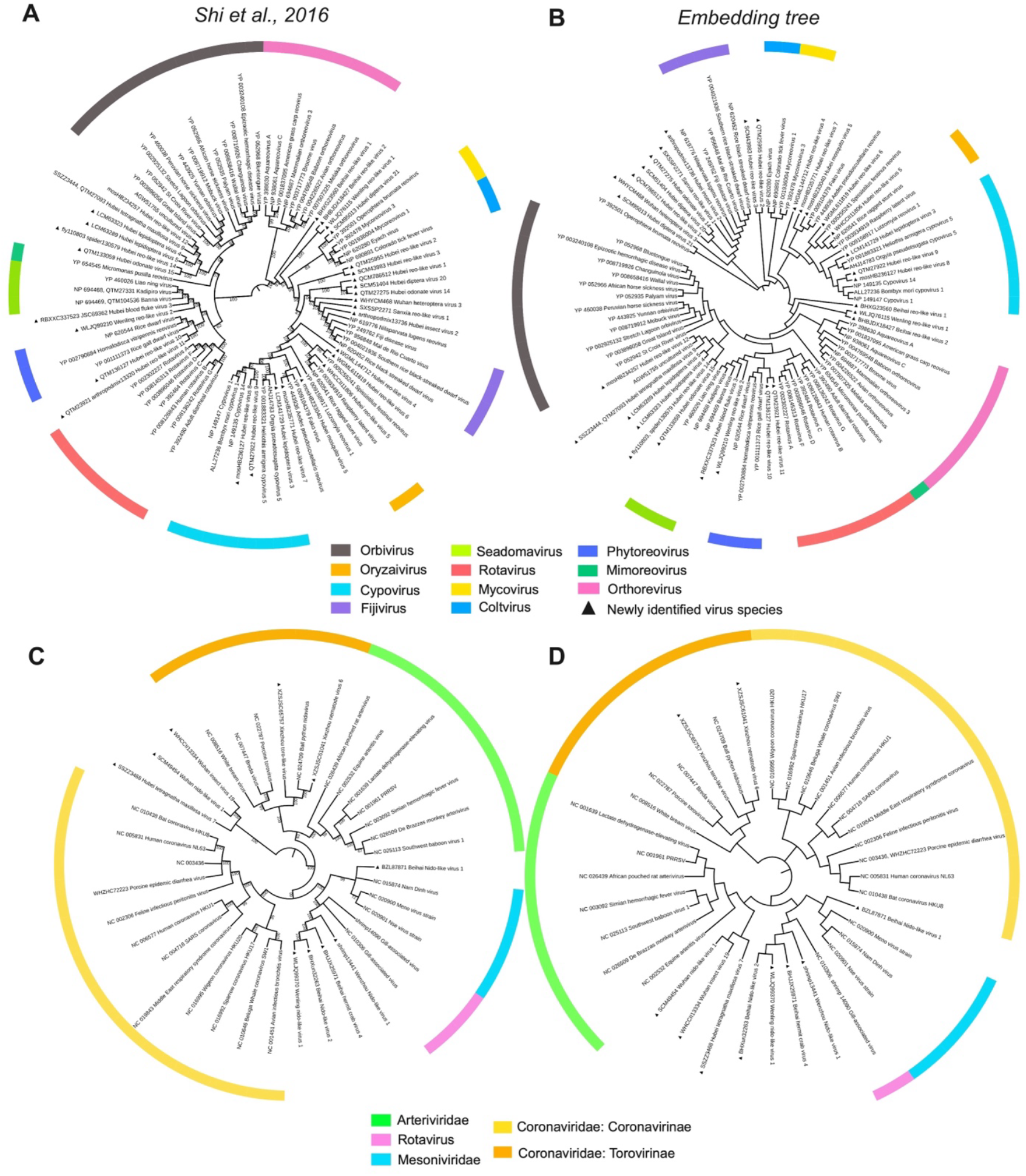
High consistency between the ML tree and embedding tree for RdRp domain. **(A-B)** The ML tree and the embedding tree constructed from layer 3 for the “Reo” clade, respectively. **(C-D)** present the same as (A-B), but for the “Nido” clade with embeddings from layer 2. For a fair comparison, the branch lengths were not displayed. The trees were visualized using iTOL web tool.^59^

In terms of broad viral classifications, the ML tree of the “Reo” clade was divided into several groups, including “Orbivirus”, “Orthorevirus”, “Rotavirus”, and others, based on current knowledge of evolutionary relationships (**Figure 4A**). The embedding tree (**Figure 4B**) also reflected these groupings. Moreover, the embedding tree not only recovered the major virus genera from the original study (based on ML) but also maintained complete consistency with the ML tree’s topology at the subgenus level, such as for “Rotavirus” and “Cypovirus”. Additionally, for newly discovered viruses, the embedding tree positioned them in a manner consistent with the ML tree.

Despite the broad agreement between the ML and embedding trees, indicated by an RF score of 0.77 and a CI score of 0.83, some differences exist. From a user’s point of view, ML inference is able to provide statistical branch support values, which are absent in the embedding tree since the extracted embeddings are fixed relative to the MSA. Moreover, there are minor topological differences in the arrangement of certain groups, which lack strong bootstrap support. For example, in the ML tree, Oryzaivirus is placed with Fijivirus (47%), whereas in the embedding tree, it is more closely related to Cypovirus.

Considering the “Nido” clade, the embedding tree (**Figure 4D**), much like the ML tree (**Figure 4C**), arranged virus species in the same group: “Arteriviridae”, “Rotavirus”, “Mesoniviridae”, “Coronaviridae: Coronavirinae”, and “Coronaviridae: Torovirinae”. Noteworthy, the embedding tree grouped Coronaviridae: Coronavirinae closer to Coronaviridae: Torovirinae, revealing a more nuanced evolutionary relationship that was not depicted in the ML tree and, more importantly, aligns better with current scientific understanding.^43^ Taken together, the embedding trees exhibited a high degree of concordance with the ML tree in representing the evolutionary relationships within the “Astro” and “Orthomyxo” clades (**Figure S22**). This consistency underpins the robustness of embedding trees in elucidating evolutionary relationships, while suggesting some avenues for further consideration.

### 2.6 The embedding tree sheds novel light on NCLDV evolution

What are the key differences between the ML tree and the embedding tree, and can the embedding tree be leveraged to address more complicated evolutionary problems? To answer those questions, we investigated the origin and evolution of NCLDVs. NCLDVs are a group of double-stranded DNA viruses known as “giant viruses” due to their exceptionally large genome size.^44^ One of the key challenges in NCLDV evolution studies is determining whether their conserved core proteins reflect a shared evolutionary history, or if some were acquired repeatedly from hosts.^45^ To study NCLDV evolution, Guglielmini et al. first identified the eight most conserved proteins and then constructed ML trees for each protein individually.^45^ They confirmed a broad concordance among the eight single-protein ML trees, indicating a shared phylogenetic signal; however, some trees displayed low bootstrap support values, and several families were not monophyletic (**Figure S23**).

We are examining whether the inconsistencies arise from the method used, or if they are inherent to the data. To this end, we built embedding trees for these proteins to assess whether the embedding trees support or contradict existing findings. The MSAs of those proteins contain a large number of gaps that Guglielmini and colleagues deemed interfere with accurate tree inference, hence trimmed them “post” application of sequence alignment; the original MSA is referred to as the “pre-trimmed” MSA. In our analysis, due to the input length limitations of the MSA Transformer, we utilized the “post-trimmed” MSAs for seven of the eight core proteins to construct embedding trees across different layers (**Table S7**). We compared the ML trees with the embedding trees for the same proteins, using the post-trimmed MSAs (“STAR Methods - RdRp and NCLDV data”). The seven core proteins used are: family B DNA polymerase (DNA pol B), D5-like primase-helicase (Primase), homologs of the Poxvirus Late Transcription Factor VLTF3 (VLTF3-like), transcription elongation factor II-S (TFIIS), genome packaging ATPase (pATPase), major capsid protein (MCP), and RNA polymerase subunit-b (RNAP-b). We observed that the embedding trees from layer 2, or layer 3, showed the greatest similarity to their ML trees (**Table S8**). Rather than relying solely on the highest RF score, we based our selections on the evolutionary relationships in the embedding trees from layers 2 or 3, given their minimal differences in RF scores.

#### The embedding tree demonstrates clearer monophyletic groupings for Ascoviridae, Mimiviridae, and the Extended mimiviridae

The ML tree of TFIIS protein was the least supported, likely due to it being the shortest marker, with several families having one or two taxa branching outside of well-recognised families (**Figure 5A**). For instance, *Diadromus pulchellus ascovirus 4a* diverges outside Ascoviridae, while *Phaeocystis globosa virus* separates from the Extended Mimiviridae, both backed by low bootstrap values (13% and 3%, respectively). The ML tree derived from the larger marker, the Primase protein, *Cafeteria roenbergensis virus* branches just outside, but near the Mimiviridae family (**Figure 5B**). However, their embedding tree demonstrates clearer monophyletic groupings, albeit without bootstrap value support. In the embedding tree for TFIIS protein, *Diadromus pulchellus ascovirus 4a* is grouped with the Ascoviridae family, and *Phaeocystis globosa virus* aligns with the Extended Mimiviridae family (**Figure 5E**). In the Primase protein embedding tree, *Cafeteria roenbergensis virus* is positioned with Mimiviridae (**Figure 5F**). These viruses consistently formed monophyletic groups across 7 embedding trees (**Figure S24**), but not in the ML tree (**Figure S23**).

**Figure 5:**
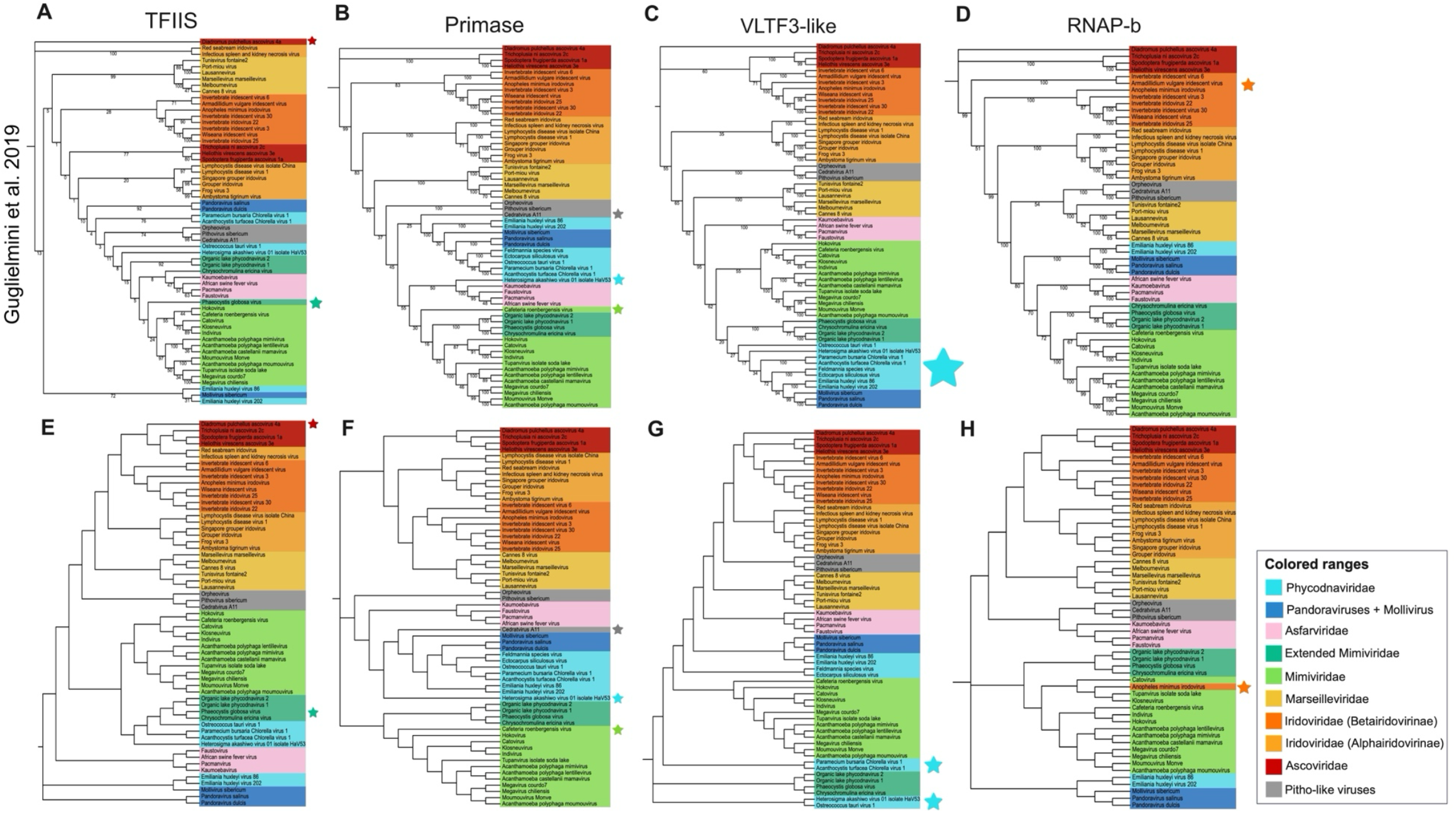
Comparison between the ML trees and embedding trees for NCLDVs. The ML trees of four core proteins for NCLDVs: **(A)** TFIIS, **(B)** Primase, **(C)** VLTF3-like, and **(D)** RNAP-b. **(E)** The TFIIS embedding tree constructed from layer 2. **(F)** The Primase embedding tree constructed from layer 3. **(G)** The VLTF3-like embedding tree constructed from layer 3. **(H)** The RNAP-b embedding tree constructed from layer 2. To enhance the reader’s understanding, the differences between the embedding tree and the ML tree discussed are highlighted with asterisks (^*^). For a fair comparison, branch lengths were omitted. The trees were visualized using iTOL web tool.^59^

#### The embedding tree appears non-monophyletic for Phycodnaviridae and Pitho-like virus

The authors suggested that single-protein ML trees indicated a shared phylogenetic signal. However, the bootstrap values for Phycodnaviridae across all ML trees were relatively low, failing to provide robust support for this inference (**Figure S23**). Intriguingly, the Phycodnaviridae exhibited completely different patterns in the ML trees and embedding trees for Primase and VLTF3-like proteins: one grouping was monophyletic, while the other was non-monophyletic (**Figures 5B-5C** and **5F-5G**). For example, in the Primase protein ML tree, the *Heterosigma akashiwo virus 01 isolate HaV53* branched outside, but close to the Phycodnaviridae family (**Figure 5B**), which was nested within this family for the embedding tree (**Figure 5F**). More importantly, the Phycodnaviridae family appeared non-monophyletic in the embedding trees for the other six proteins (**Figure S24**). Moreover, the monophyletic grouping of Pitho-like viruses does not hold in the ML tree (**Figure 5B**), which is more apparent in the embedding tree of the Primase protein (**Figure 5F**).

#### The MSA Transformer is sensitive to gaps in the MSA

Both the ML tree and the embedding tree indicated a monophyletic relationship for RNAP-b protein (**Figures 5D** and **5H**), except for *Anopheles minimus iridovirus*. In the embedding tree, *Anopheles minimus iridovirus* branches outside Betairidovirinae but is grouped with Mimiviridae (**Figure 5H**). Upon closer examination, we observed that in the post-trimmed MSA, the sequence of *Anopheles minimus iridovirus* contained extensive gaps covering approximately 300 amino acids. After removing the long-gap region from the whole MSA of the RNAP-b protein, we reconstructed the embedding tree using the same layer, which resulted in both trees exhibiting the same monophyletic relationship (**Figure S24H**).

#### MSA Transformer does not consistently outperform traditional ML methods when handling gappy alignments

To better assess the capability of both the MSA Transformer and the ML approach in recovering phylogenetic relationships from gappy MSAs, we conducted four comparative analyses: (1) comparing the embedding tree inferred from a length-controlled “gappy” MSA (a shortened version of the gappier, pre-trimmed MSA) with the ML tree constructed from the post-trimmed MSA; the gappy MSA contained the same number of columns as the post-trimmed, which is a random sub-set of the pre-trimmed MSA, (2) comparing the embedding tree from a gappy MSA with the embedding tree from the post-trimmed MSA, (3) comparing the ML tree from the gappy MSA with the ML tree from the post-trimmed MSA, and (4) comparing the ML tree from the gappy MSA with the embedding tree from the post-trimmed MSA. The performance of each method was evaluated by calculating the average CI scores between trees inferred from gappy and post-trimmed MSAs, across 50 replicates for each of the four comparisons (see “STAR Methods - Phylogenetic tree construction from gappy MSAs”). Our findings indicate that the MSA Transformer does not consistently outperform the ML method in reconstructing phylogenetic relationships established from the curated MSAs, from the gappier versions, particularly in certain protein families such as RNAP-b and DNA pol B (**Figure S25**).

As evolution cannot be observed directly, it is impractical to determine if traditional or language model-based reconstruction methods are more accurate. The embedding trees, however, provide a different explanation for where the low bootstrap values are placed. Our embedding trees do not support the conclusion that conserved proteins in NCLDVs share a common evolutionary history but instead suggest independent evolution.

## 3. Discussion

### The MSA Transformer does not recapitulate classical phylogenetics

Classical phylogenetic inference methods typically assume that sites in a sequence evolves independently. We propose that MSA Transformer may address this limitation, as it has the capability to automatically learn the relationships both between sequences and between amino acids. We sought to substantiate this claim with several lines of supporting evidence. When comparing embedding trees from the original MSAs to those from shuffled-columns MSAs, we observe a progressive decrease in similarity across layers (**Figures 3A**). For instance, in PF13377, there is 100% concordance between trees generated from the original and shuffled-columns MSA in the first layer, highlighting column conservation; meanwhile, the later layers indicate the model’s ability to capture sequence context. Besides, synthetic embedding trees showed significantly higher similarity to their corresponding ML trees than those from Pfam MSAs (**Figure 3B**). The high similarity may be attributed to the absence of epistatic effects in the synthetic MSAs; this absence enables MSA Transformer and likelihood-based methods to confirm the same zero-order relationships from the MSAs. More importantly, we implemented a ridge model to predict the epistatic effects of avGFP double mutants, based on concatenated embeddings from the MSA Transformer. We experimented with different random seeds and found that achieving statistical significance required careful tuning of the model’s hyperparameters. Although this may be attributable to the simplicity of the model or the limited sample size, these results nonetheless indicate that MSA Transformer implicitly encodes information relevant to epistasis.

Unlike traditional methods that generally overlook gaps in sequence alignments,^46^ the MSA Transformer takes these gaps into account. To determine whether this feature enhances the accuracy of evolutionary analysis, we compared the performance of the MSA Transformer with that of a likelihood-based approach using alignments containing many gaps from seven NCLDV protein families. The MSA Transformer did not consistently perform better than the likelihood-based method; in several cases, including RNAP-b and DNA pol B, the likelihood-based method actually produced results more similar to those obtained from higher-quality, less gappy data. These findings suggest that simply considering gaps in the analysis does not necessarily lead to improved evolutionary inference, regardless of how many gaps are present in the alignment.

### There are limitations in applying MSA Transformer to phylogenetic research

First, alignment-based approaches have shown constraints in scenarios where MSA quality is low.^47,48^ This issue also impacts MSA Transformer, which depends on information within columns to infer phylogenetic relationships. Second, different layers of embeddings vary in their ability to recover the evolutionary history of proteins. We do not have an explicit guide for which layer is optimal in a specific case, but we recommend using early layers 2 to 5, as they most closely resemble traditional phylogenetic trees. Third, although MSA Transformer can theoretically handle up to 1,024 protein sequences with an average alignment length of 1,022, its practical capacity is limited by memory requirements, making it less suitable for large-scale phylogenetic analyses.

### Future directions

Transformer-based language models act as advanced compression systems, processing sequences to generate rich representations that are highly effective for downstream tasks.^49^ Our study compares an MSA-based language model with alignment-based phylogenetic reconstruction methods. We found embedding trees derived from both the original MSA and the shuffled-column MSA exhibit comparable similarity to traditional phylogenetic trees across all layers, while the deeper layers show increased sensitivity to the biological authenticity of sequences. This suggests that the current positional encoding mechanism may not strongly influence model performance, especially in early layers. Building on this observation, we aim to incorporate bootstrap support into late-layer embedding trees by shuffling the columns of the original MSA. In addition, MSA Transformer supports phylogenetic reconstruction only for protein sequences, underscoring the need to train models that can operate at the genomic level and manage longer sequences.^50^ We anticipate that PLMs will complement classical phylogenetic methods, contributing to more accurate reconstructions of protein family evolutionary histories.^40,51^

## ACKNOWLEDGMENTS

This work was supported by UQ Graduate School Scholarships to R.C.

## AUTHOR CONTRIBUTIONS

All authors designed the study. R.C. implemented the code, performed the computational experiments, and wrote the manuscript. M.B. supervised the project. All authors contributed to reviewing and editing the final paper.

## STAR★METHODS

Detailed methods are provided in the online version of this paper and include the following:

### KEY RESOURCES TABLE

| REAGENT or RESOURCE | SOURCE | IDENTIFIER |
| --- | --- | --- |
| Deposited data |  |  |
| Simulated MSAs | This study | https://doi.org/10.5281/zenodo.16923353 |
| ML tree for Pfam MSAs | This study | https://doi.org/10.5281/zenodo.16923353 |
| Pfam MSAs and NJ trees | Pfam version 35.0,<br>November 2021 | https://www.ebi.ac.uk/interpro/entry/pfam |
| RdRp domains | Shi et al.<sup>42</sup> | https://www.nature.com/articles/nature20167 |
| NCLDV domains | Guglielmini et al.<sup>45</sup> | https://www.pnas.org/doi/10.1073/pnas.1912006116 |
| avGFP DMS dataset | Chen et al.<sup>41</sup> | https://doi.org/10.1016/j.cels.2023.07.003 |
| Software and algorithms |  |  |
| Code and scripts | This study | https://github.com/Cassie818/MsaPhylo and<br>https://doi.org/10.5281/zenodo.16923353 |
| IQ-TREE version 2.1.2 | Nguyen et al.<sup>15</sup> | http://www.iqtree.org/ |
| GRASP version 29-Nov-2023 | Foley et al.<sup>52</sup> | http://grasp.scmb.uq.edu.au/ |
| MAFFT version 7 | Katoh et al.<sup>53</sup> | https://mafft.cbrc.jp/alignment/software/ |
| MSA Transformer | Rao et al.<sup>22</sup> | https://github.com/facebookresearch/esm |
| ETE version 3.1.3 | Huerta-Cepas et al.<sup>54</sup> | https://etetoolkit.org/ |
| SciPy version 1.11.4 | Virtanen et al.<sup>55</sup> | https://scipy.org/ |
| NumPy version 1.25.2 | Harris et al.<sup>56</sup> | https://numpy.org/ |
| Matplotlib version 7.1 | Hunter et al.<sup>57</sup> | https://matplotlib.org/ |
| TreeDist version 2.6.2 | Smith et al.<sup>38</sup> | https://github.com/ms609/TreeDist |
| Scikit-learn version 1.6.1 | Pedregosa et al.<sup>58</sup> | https://scikit-learn.org/stable/ |
| NJ algorithm | Yeung et al.<sup>35</sup> | https://github.com/esbgkannan/chumby |
| iToL version 6 | Letunic et al.<sup>59</sup> | https://itol.embl.de/ |

### RESOURCE AVAILABILITY

#### Lead contact

Further information and requests for resources and reagents should be directed to and will be fulfilled by the lead contact, Mikael Bodén.

## Materials availability

No new materials were generated in this study.

## Data and code availability

- This paper analyzes existing, publicly available data. These accession numbers for the datasets are listed in the key resources table.
- All original code has been deposited at GitHub repository (https://github.com/Cassie818/MsaPhylo) and is publicly available as of the date of publication. DOIs are listed in the key resources table.
- Any additional information required to reanalyze the data reported in this paper is available from the lead contact upon request.

## METHOD DETAILS

### Pfam data

Pfam is a comprehensive database that includes Pfam-A, a manually curated collection of protein families. These families are based on significant sequence similarity, indicating homology.^60^ Each Pfam-A entry includes a seed alignment, a representative set of sequences that serve as the foundation for constructing profile hidden markov models with HMMER software.^61^ The utility of seed alignments from Pfam-A enables efficient and sustainable manual curation of both alignments and annotations.^62^

In our study, twenty unique Pfam-A seed alignments, along with their NJ trees generated using FastTree^63^ software, were downloaded from the Pfam^64^ database (version 35.0, November 2021). We utilised IQ-TREE^15^ to construct the related ML trees with the Le Gascuel (LG) substitution model. All parameters used their default values unless specified otherwise.

We conducted statistical analyses on collected MSAs, NJ trees, and self-constructed ML trees across 20 Pfam protein families. An MSA is defined as a matrix *X* ∈ ℝ^*M*×*L*^, where *M* represents the number of sequences (rows), and *M* indicates alignment lengths (columns). Each matrix entry *x*_*mj*_ is encoded as an integer value, signifying the specific amino acid identity at position *j* in sequence *m* .^22^ We calculated the mean sequence lengths *µ*_*L*_, excluding gaps, to estimate the integrity and utility of the alignments.

Furthermore, we computed the mean evolutionary distances *δ*, for both NJ and ML trees, reflecting phylogenetic depth and measuring evolutionary divergence among the sequences analyzed. Let *T* be a tree with a set of leaf nodes {*v*_1_, *v*_2_, …, *v*_*M*_}. Then, for each leaf node *v*_*i*_, *path*(*v*_*i*_, *root*) denote the sequence of nodes from leaf *v*_*i*_ to the root of the tree, and let *dist*(*e*) denote the distance from node *e* to its parent node. The distance from a leaf node *v*_*i*_ to the root is calculated by summing the distances along the path from the leaf to the root:

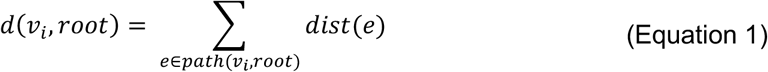

Thus, the mean evolutionary distances of the tree can be defined as the average of these distances for all leaf nodes:

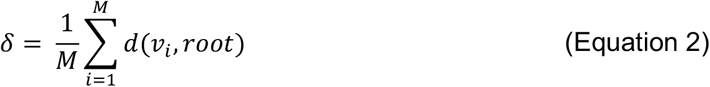

### Synthetic MSAs

For each Pfam protein family, we employed TrAVIS, a component of the GRASP^52^ suite (version 29-Nov-2023), to simulate MSA. TrAVIS samples branch lengths from a gamma distribution *γ*(*κ, θ*), where *κ* denotes the shape parameter and *θ* represents the scale parameter. These distances are subsequently normalized to the mean evolutionary distances *δ*. The tree generation process iteratively bifurcates each branch point until the specified number of sequences has been mapped as leaves.

We first set the *extants* parameter to match the number of sequences *M* in the original MSA. To ensure the synthetic MSA had a similar amino acid composition as the original but lacked biological meaning, we randomly selected a protein sequence from the shuffled-columns MSA and removed the gap character “-”. Then, the evolution simulation started at the root with the selected protein sequence under the LG substitution model, determining the sequence at each child recursively. For a sequence, each position is considered as a possible site for a mutation as a Poisson process with a relative rate *r* .

After generating the synthetic sequences, we aligned them into an MSA using MAFFT^53^ and constructed the ML tree using IQ-TREE^15^, following the same method used for the Pfam MSA. This process was repeated multiple times to adjust the parameters *rates* and *dist*, ensuring that the synthetic MSA closely approximated the alignment lengths *L* of the original MSA and that its ML tree had a similar mean evolutionary distance *δ* to the Pfam ML tree. The parameters employed by TrAVIS are detailed in **Table S3**, while **Table S4** presents a comparison between the original MSA and synthetic MSA for each Pfam protein family.

### RdRp and NCLDV data

The MSAs and ML trees for the RNA virus RdRp domain were directly sourced from the work of Shi et al.^42^ In addition, the pre-trimmed MSAs, post-trimmed MSAs, and ML trees related to NCLDVs were obtained from the study by Guglielmini et al.^45^ Post-trimmed MSAs were generated by removing columns with more than 20% gaps from the pre-trimmed alignments.

Statistical details regarding the collected MSAs and ML trees are summarized in **Tables S5 and S7**. Detailed methodologies for phylogenetic tree construction can be found in the Methods sections of these two referenced papers.

### Phylogenetic tree construction from gappy MSAs

For each NCLDV protein family, we generated “gappy MSAs” by randomly sampling columns from the pre-trimmed MSA, resulting in alignments with the same number of columns as the post-trimmed MSA but containing more gaps. This sampling procedure was repeated 50 times to ensure statistical robustness. For each replicate, a ML tree was reconstructed from the gappy MSA using IQ-TREE^15^ with the LG model, and an embedding tree was generated using the same method described above. For both gappy and post-trimmed MSAs, embedding trees were constructed strictly using embeddings at the same layer.

### Epistasis calculation and epistatic interactions identification

We obtained the DMS dataset for single and double variants of the avGFP protein from Chen et al.^41^ Epistasis (*ε*_*ij*_) for the pair of mutations *i* and *j* is calculated as the difference between the observed log-fitness of the double mutant and the sum of the log relative fitness of the single mutants, that is: ^65^

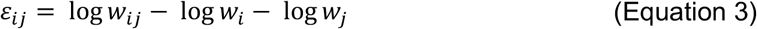

where log *w*_*ij*_ denotes the log relative fitness of the double mutant, and log *w*_*i*_ and log *w*_*j*_ represent the logarithms of the fitness values of the individual single mutants, respectively.

Double variants with absolute epistasis values exceeding one standard deviation across 1,000 selected samples were classified as epistatic variants, while those within one standard deviation were considered non-epistatic.

### MSA Transformer, embeddings, and column attention

We employed MSA Transformer,^22^ namely the ESM-MSA-1b model with 100 million parameters, and it was implemented via the fair-esm package. Unlike other PLMs, MSA Transformer is designed to process multiple aligned protein sequences with *M* rows and *L* columns. A start token is added to the input MSA, and each amino acid or gap character “-” is encoded as a vector in ℝ^768^.

The model was constructed with 12 layers, each comprising a 12-head row attention block, a 12-head column attention block, and a feed-forward network. At each layer *l* (*l* = 1, …,12), it produces embeddings represented as a three-dimensional array *X*^(*l*)^ ∈ ℝ^*M* × (*L* + 1) ×*d*^, where *L* + 1 corresponds to the alignment length inclusive of the special start token. Each column *j* of this array is represented as 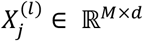. To obtain column attention, the MSA Transformer first employs three specific *d* × *d* matrices: 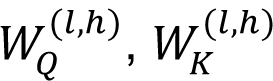, and 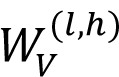, to create the “query,” “key,” and “value” vectors, respectively. Then, for each layer *l* and head *h* (both ranging from 1 to 12), three matrices in ℝ^*M* ×*d*^ were computed for each column *j* through the following transformations:

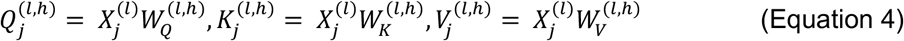

The column attention head for each column *j*, at layer *l*, and from head *h*, is therefore derived as follows:

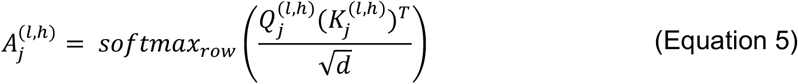

where *softmax*_*row*_ applies the softmax function to each row of a matrix independently, converting the elements of each row into a probability distribution.

### Spearman rank correlation coefficients with evolutionary distances

We removed the start token from the embedding and column attention across all protein families, as it lacks substantive meaning. For each layer *l*, the pairwise Euclidean distances of the three-dimensional embeddings matrix were computed, resulting in a *M* × *M* distance matrix *D*^(*l*)^. More specifically, the Euclidean distance between the *p*-th and *q*-th sequences (or rows) of the matrix *X*^(*l*)^ from layer *l*, denoted as 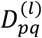, is defined by the following equation:

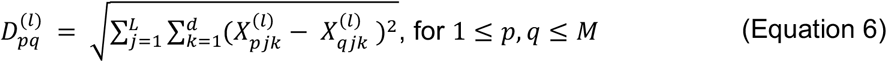

where 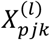 and 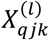 represent the *j*-th column and *k*-th dimension of embeddings at layer *l*, corresponding to the *p*-th and *q*-th sequences, respectively.

Previous studies demonstrated that some CMSCAHs capture the Hamming distances.^22,31^ Building on this, we also averaged the *M* × *M* column attention head matrices 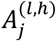 from Equation (5) across all columns of the MSA. Subsequently, we symmetrized these matrices to obtain a *M* × *M* matrix, as:

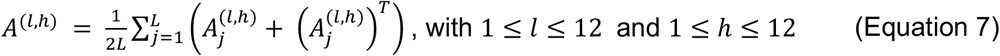

Using the ETE3^54^ toolkit, we computed the pairwise evolutionary distance matrix *E*, where *E* ∈ ℝ^*M* × *M*^, based on ML and NJ trees, separately. The evolutionary distance *E*_*pq*_ between any two nodes *p* and *q* was calculated by summing the branch lengths between them. We then computed the Spearman’s rho between the pairwise evolutionary distances matrix *E* and both the Euclidean distances matrix *X*^(*i*)^ and the column attention head matrices *A*^(*l,h*)^ for each layer and head, separately. The Spearman’s rho denoted by *ρ*, was computed using the *spearmanr* function from the *scipy*.*stats* module in the SciPy.^55^ This coefficient ranges between -1 and +1, indicating perfect negative and positive correlations respectively, with *ρ* = 0 signifying no correlation, as:^66^

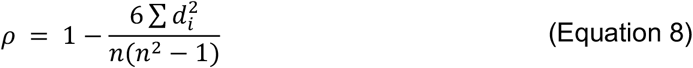

where *d*_*i*_ is the difference between the two ranks, and *n* is the number of observations.

### Phylogenetic tree reconstruction and similarity comparison

For each layer and head, we applied the NJ algorithm^4,35^ to the pre-defined Euclidean distance matrix *D*^(*l*)^ and the attention head *A*^(*l,h*)^ to construct the corresponding “embedding trees” and “attention trees.”

The TreeDist^38^ package was utilized to estimate the similarity between two phylogenetic trees, focusing on both RF and CI distances. The RF distance, commonly employed in phylogenetic research, quantifies dissimilarity between trees by counting the unique splits (bipartitions of taxa) present in one tree but absent in the other.^37^ However, it treated all non-identical splits uniformly, overlooking subtle variations and potentially introducing biases in evaluating phylogenetic trees. To fill in this gap, Smith introduced the CI distance, a more forgiving method that aligns splits from phylogenetic trees into subtrees. This approach calculated a mutual information score using the arrangement of leaf labels, effectively quantifying tree similarity by estimating the consistency in leaf grouping.^38,67^ We normalized both distances by rescaling them against a maximum value derived from the involved tree sizes and topology. Additionally, these distances were converted into similarity scores by specifying the parameter *similarity = TRUE*, termed the “RF score” and “CI score,” respectively. In this scale, a score of zero indicates complete dissimilarity, while a maximum score of one signifies identical tree structures, with higher values indicating greater similarity.

### Epistasis prediction with MSA Transformer embeddings

The MSA for avGFP protein was sourced from Chen et al.^41^ and is referred to as the wild-type MSA. Double mutations were introduced by simultaneously applying both amino acid substitutions to the first reference sequence within the MSA, while all other sequences remained unchanged; this modified MSA is referred to as the mutated MSA. Consequently, the resulting mutated MSAs contain double mutations embedded within the evolutionary context.

We randomly selected 1,000 double variants from the avGFP DMS dataset. Using the corresponding wild-type and mutated MSAs, we extracted the final layer embeddings from MSA Transformer. For a single mutation, the embedding at the mutated position in the reference sequence was obtained, yielding a vector *x* ∈ ℝ^768^. Then, for each double mutation, embeddings at the mutated positions were extracted from both the wild-type (*x*_*wt*_ ∈ ℝ^1536^) and mutated MSAs (*x*_*mut*_ ∈ ℝ^1536^). These embeddings were concatenated to form a final representation *x* ∈ ℝ^3072^ . A ridge model was trained based on the concatenated embeddings to predict epistasis values, using 10-fold cross-validation via Scikit-learn package.^58^ Model performance was assessed by computing the Pearson correlation between epistasis values measured by DMS experiments and those predicted by the model.

## Statistical analysis

Statistical tests performed by SciPy included Welch’s two-tailed and Welch’s one-tailed t-test. For shuffled MSAs, data are presented as mean ± standard deviation of embeddings. Pearson correlations were computed using the *pearsonr* function from the *scipy*.*stats* module in SciPy.^55^

## Data visualisation and analyses

We conducted analyses and created plots using R 4.3.0 and Python 3.10.12, employing the following packages: ape 5.7-1, matplotlib 7.1, numpy 1.25.2, and pandas 2.0.3.

