## Supplemental information for "Learning the Language of Phylogeny with MSA Transformer"

### Supplemental figures and figure captions

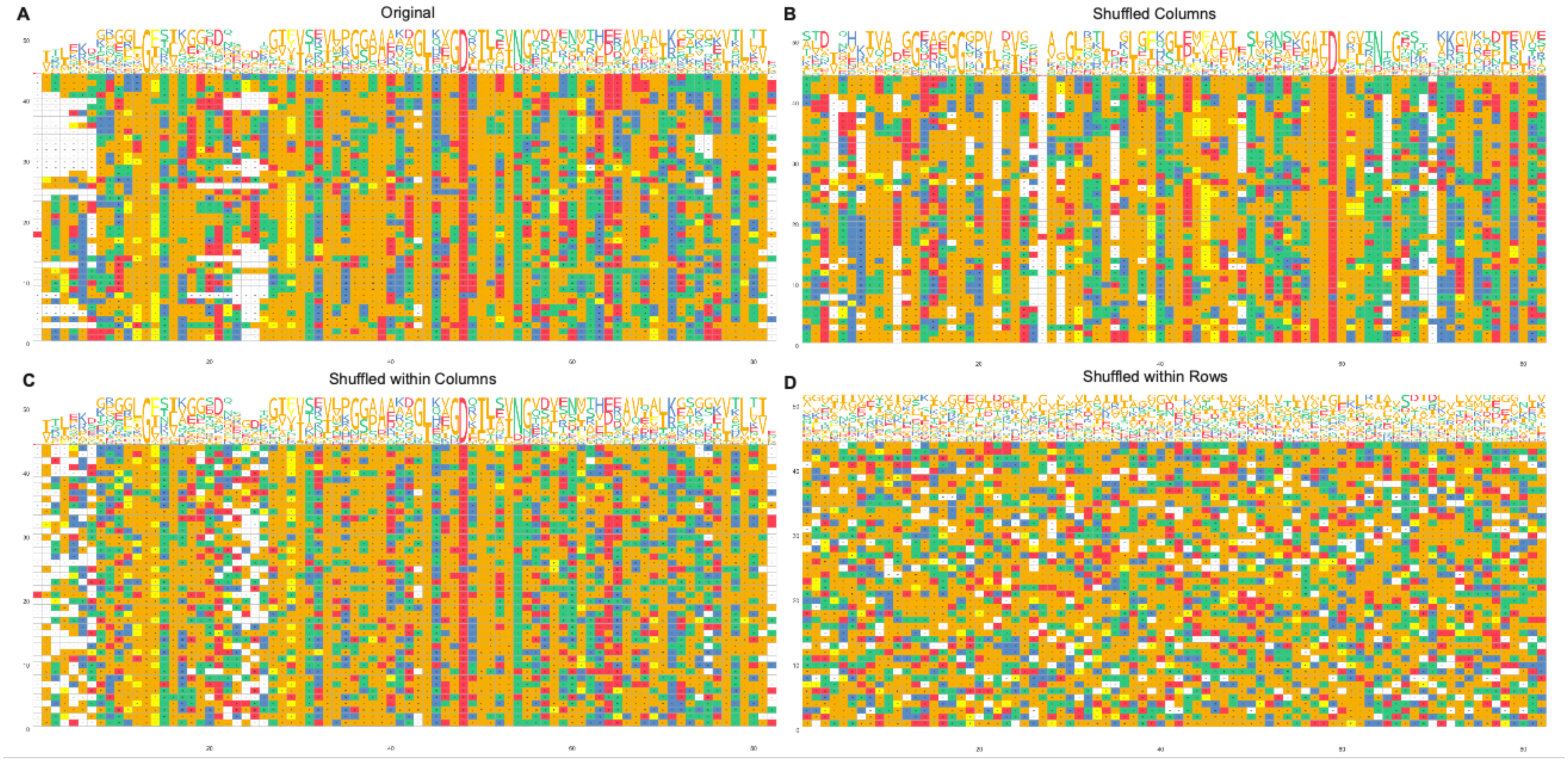


**Figure S1: Four distinct MSAs used in this study, related to Figure 1.**
(**A**) the original MSA, (**B**) the shuffled-columns MSA, (**C**) the shuffled-within-columns MSA, and (**D**) the shuffled-within-rows MSA. These alignments were plotted by ggmsa version 1.3.4.[1]

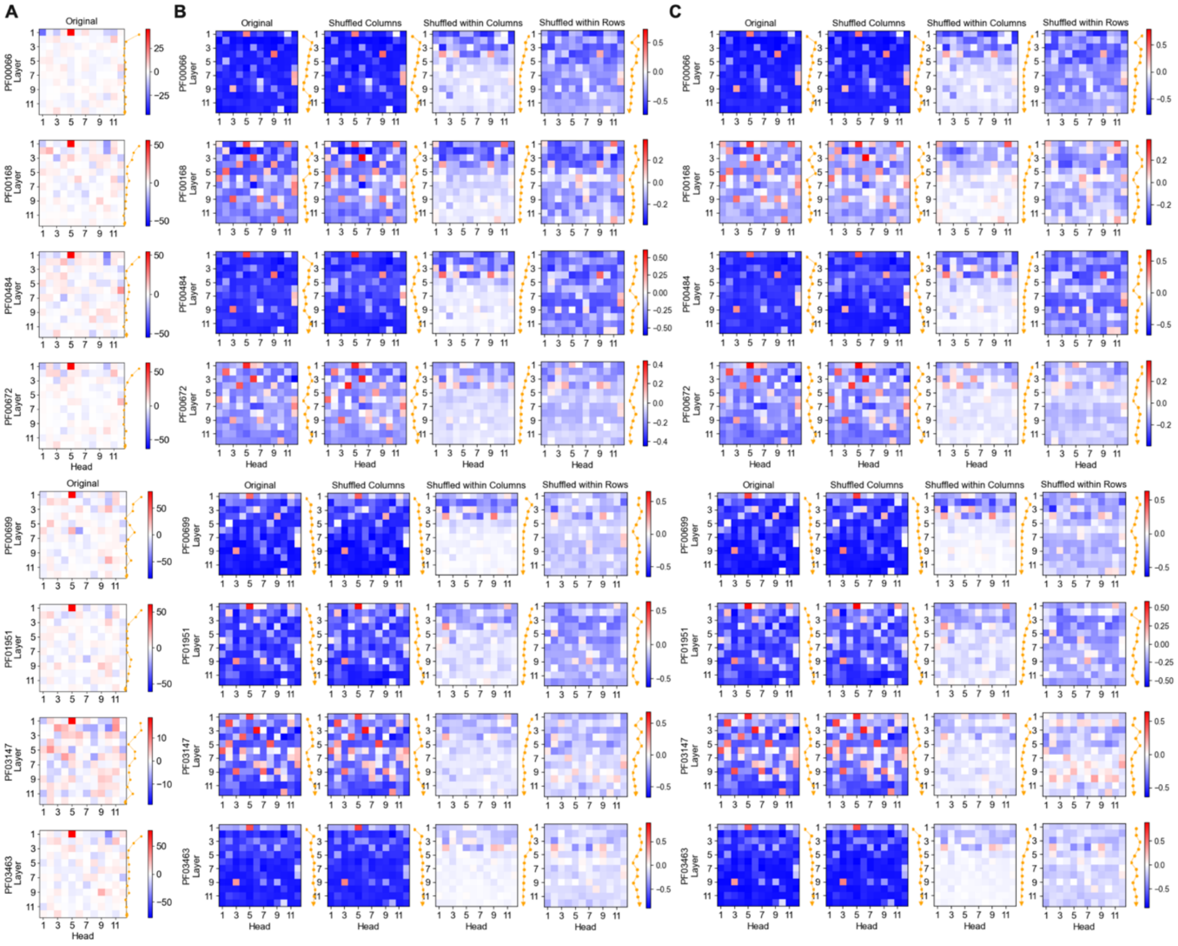


**Figure S2: The distance captured by 144 distinct column attention heads for *the previous 8* Pfam protein families, related to Figure 2.**

(**A**) Heatmap of regression coefficients from a logistic model combining 144 CMSCAHs to predict pairwise Hamming distances in the original MSA, following Lupo et al.[2]

(**B, C**) Heatmaps of Spearman's rho values between attention heads and pairwise evolutionary distances calculated from the (B) NJ tree or (C) ML tree. For shuffled MSAs, color gradients indicate the average value over five iterations (red: positive, white: zero, blue: negative). The yellow line shows layer-wise averages of absolute values across heads.

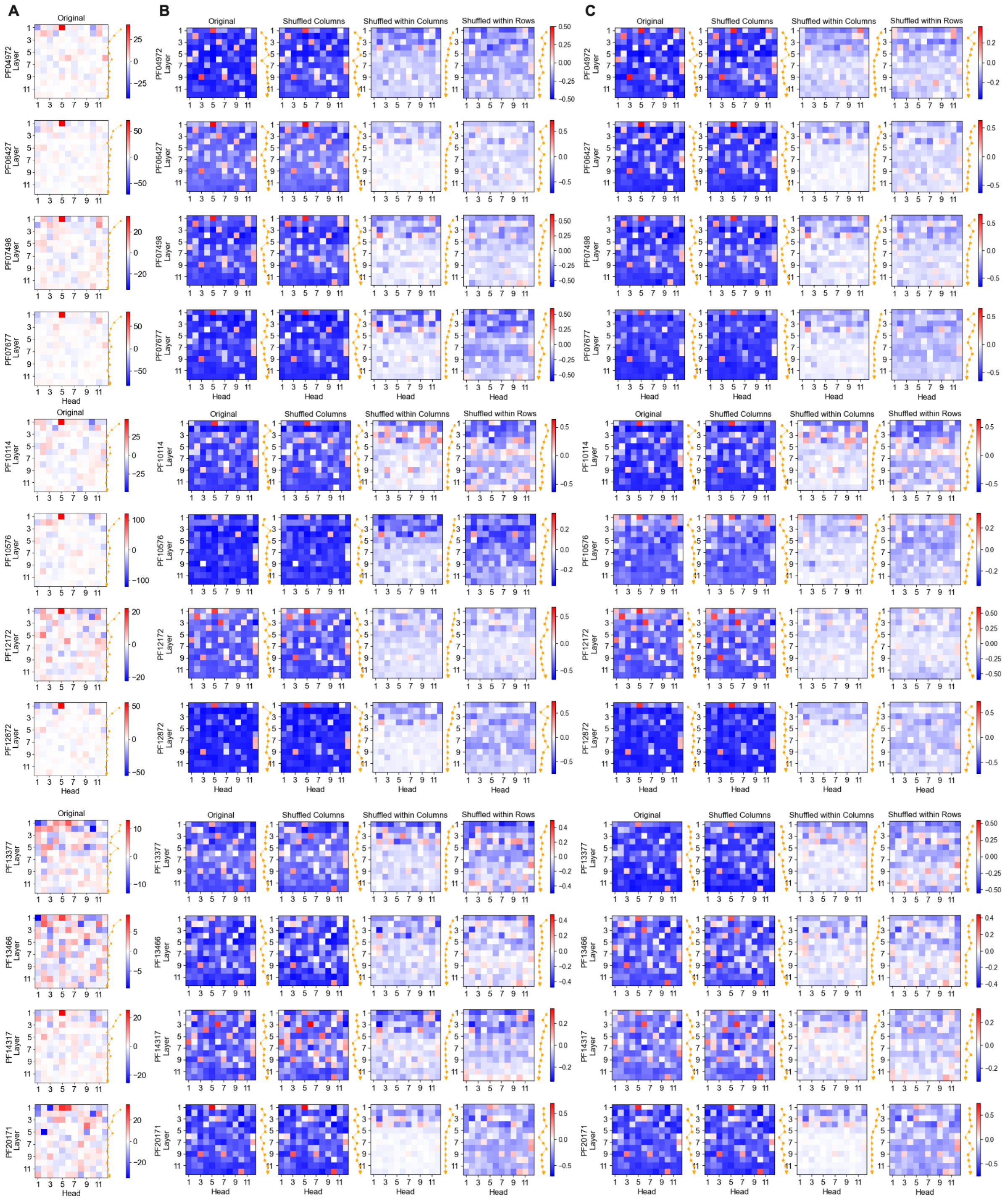

**Figure S3: The distance captured by 144 distinct column attention heads for *the rest 12* Pfam protein families, related to Figure 2.**

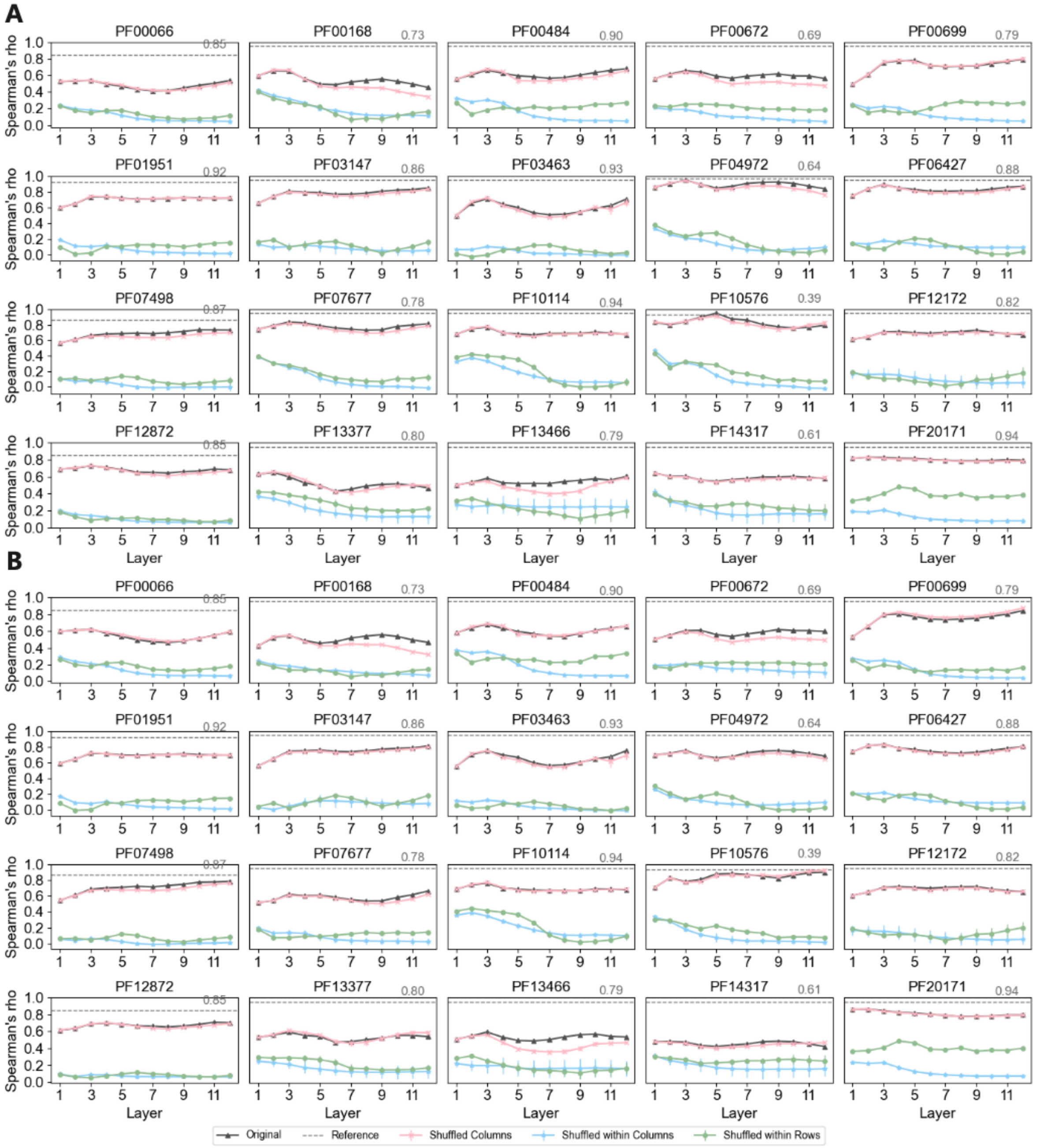


**Figure S4: The Spearman's rho between pairwise Euclidean distances of embeddings and evolutionary distances computed from two phylogenetic trees, related to Figure 2.**

(**A, B**) The pairwise evolutionary distances derived from the NJ tree and ML tree, separately. The gray reference line represents Spearman's rho between pairwise evolutionary distances derived from the NJ or ML tree for each protein family. Different types of MSAs are color-coded for clarity. For shuffled MSAs, the y-axis displays the average Spearman's rho from five shuffling iterations, with error bars indicating the standard deviation across these iterations.

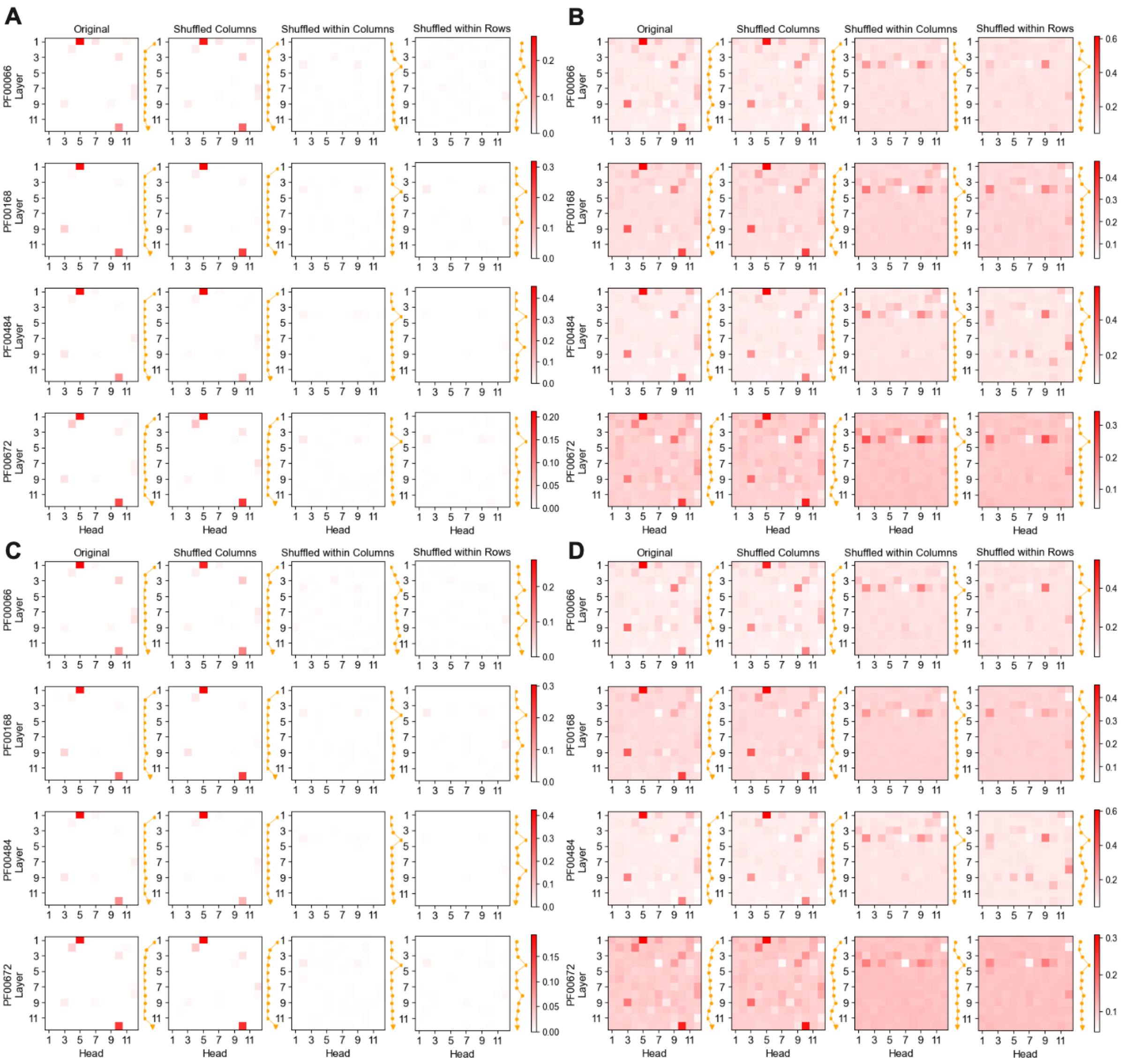


**Figure S5: The heatmap of tree similarity between attention trees and two phylogenetic trees for Pfam protein family *PF0666, PF00168, PF00484, and PF00672*, related to Figure 2.**

(**A, B**) The similarity between attention trees and *NJ* trees, measured by RF and CI scores, respectively.
(**C, D**) The similarity between attention trees and *ML* trees, measured by RF and CI scores, respectively.
For shuffled MSAs, colors indicate average similarity scores over five iterations (red: positive values, white: zero). The yellow line shows the layer-wise average of absolute values across all heads.

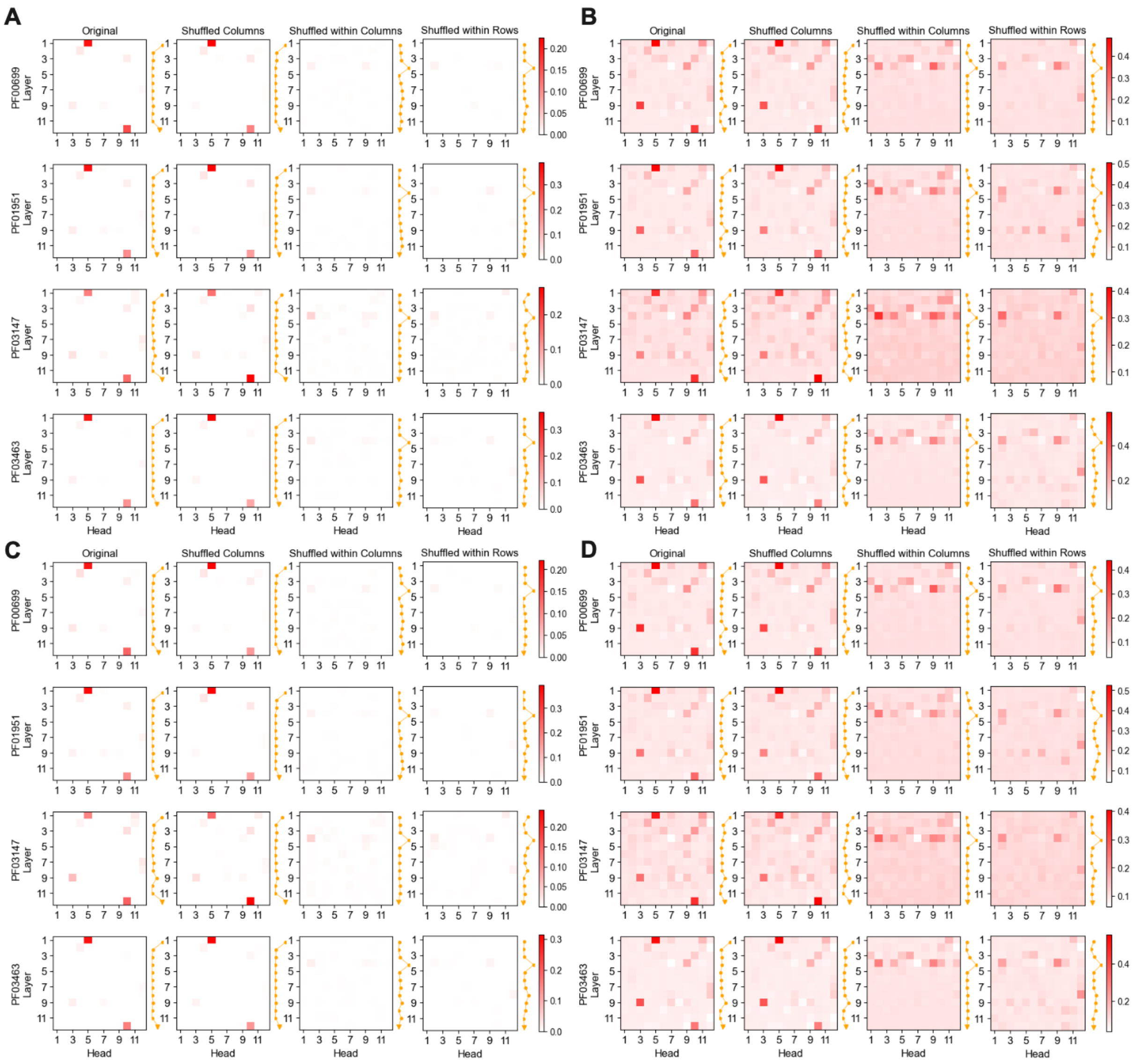


**Figure S6: The heatmap of tree similarity between attention trees and two phylogenetic trees for Pfam protein family *PF00699, PF01951, PF03147, and PF03463*, related to Figure 2.**

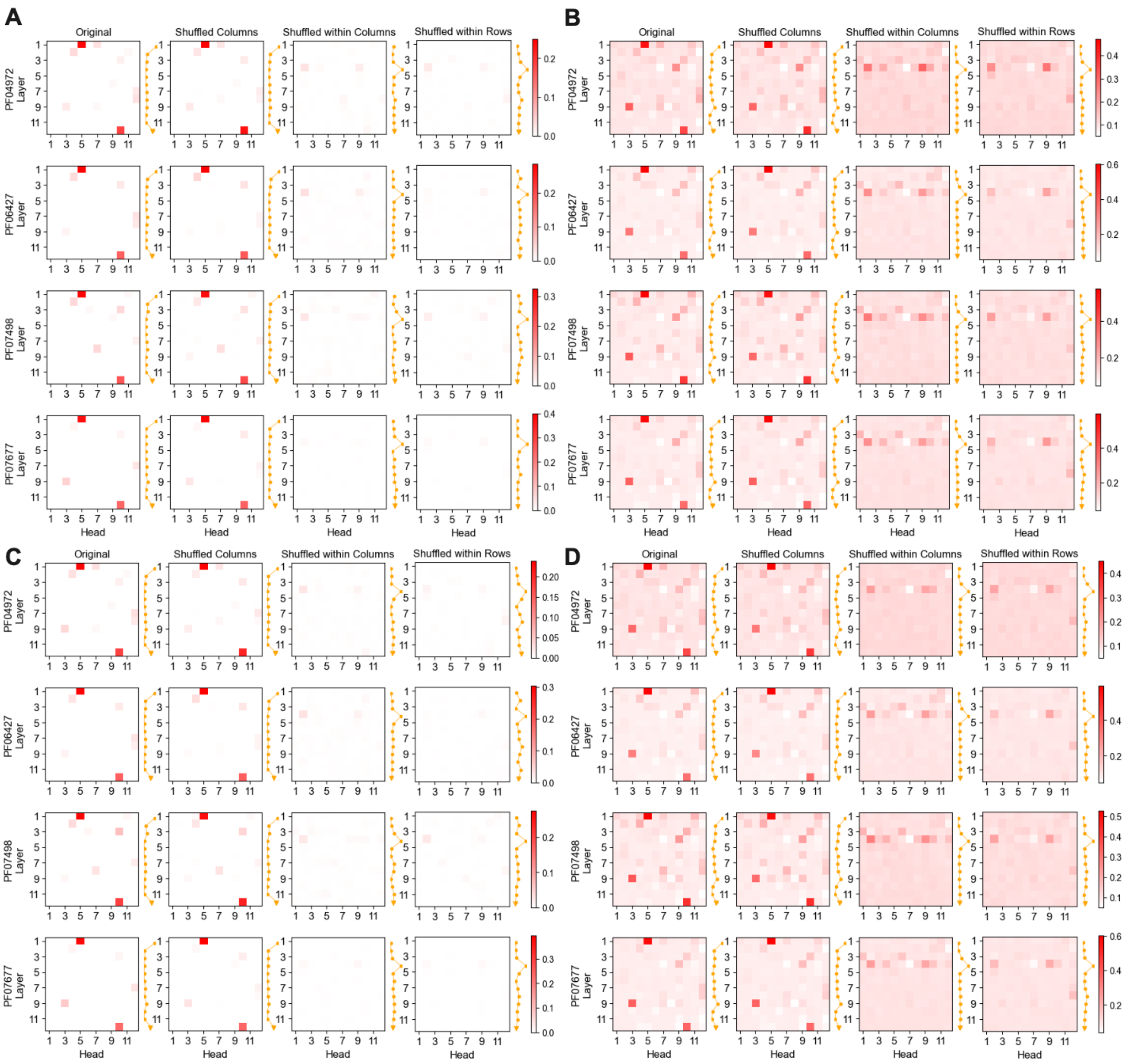


**Figure S7: The heatmap of tree similarity between attention trees and two phylogenetic trees for Pfam protein family *PF04972, PF06427, PF07498, and PF07677*, related to Figure 2.**

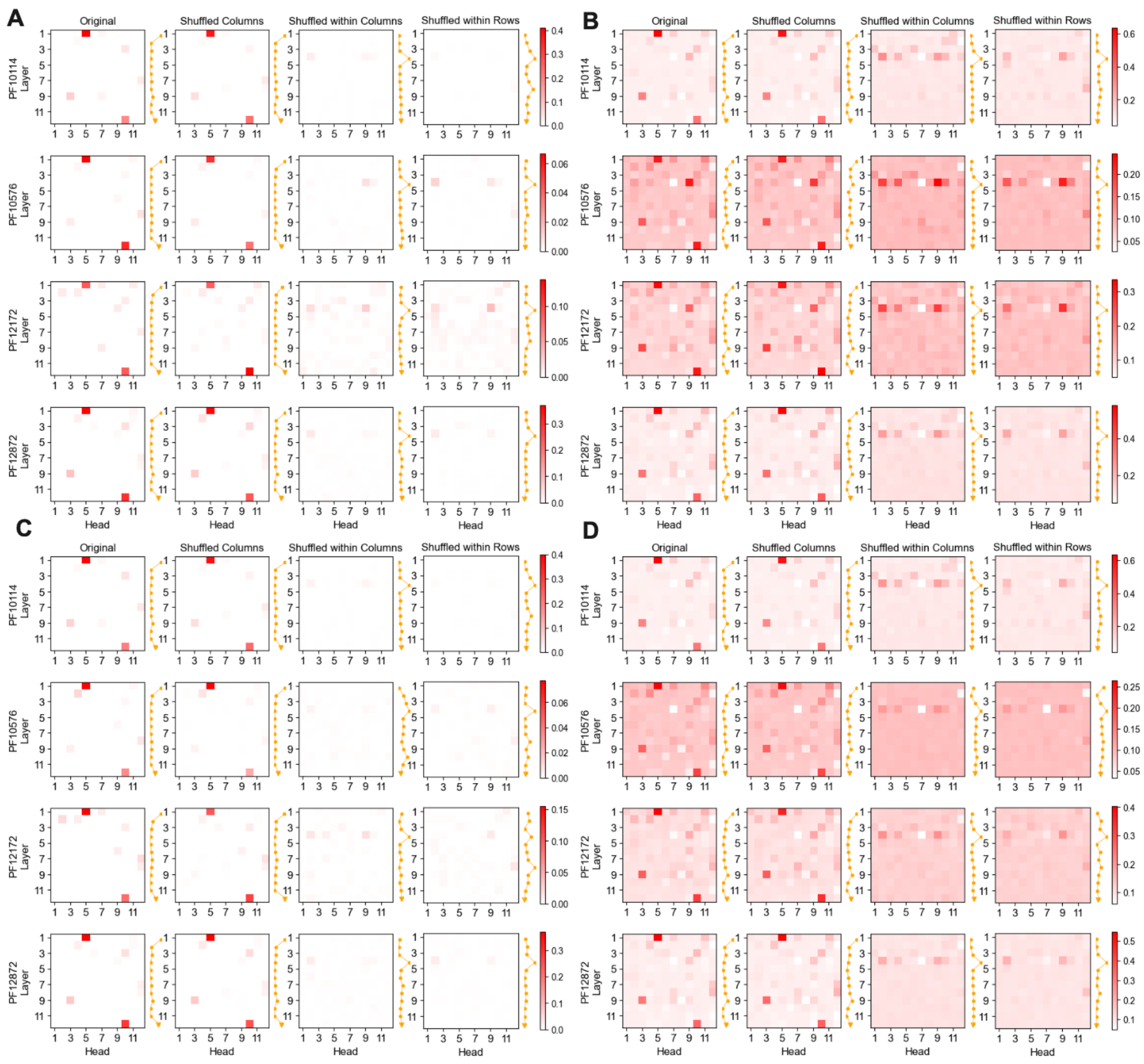

**Figure S8: The heatmap of tree similarity between attention trees and two phylogenetic trees for Pfam protein family *PF10114, PF10576, PF12172, and PF12872*, related to Figure 2.**

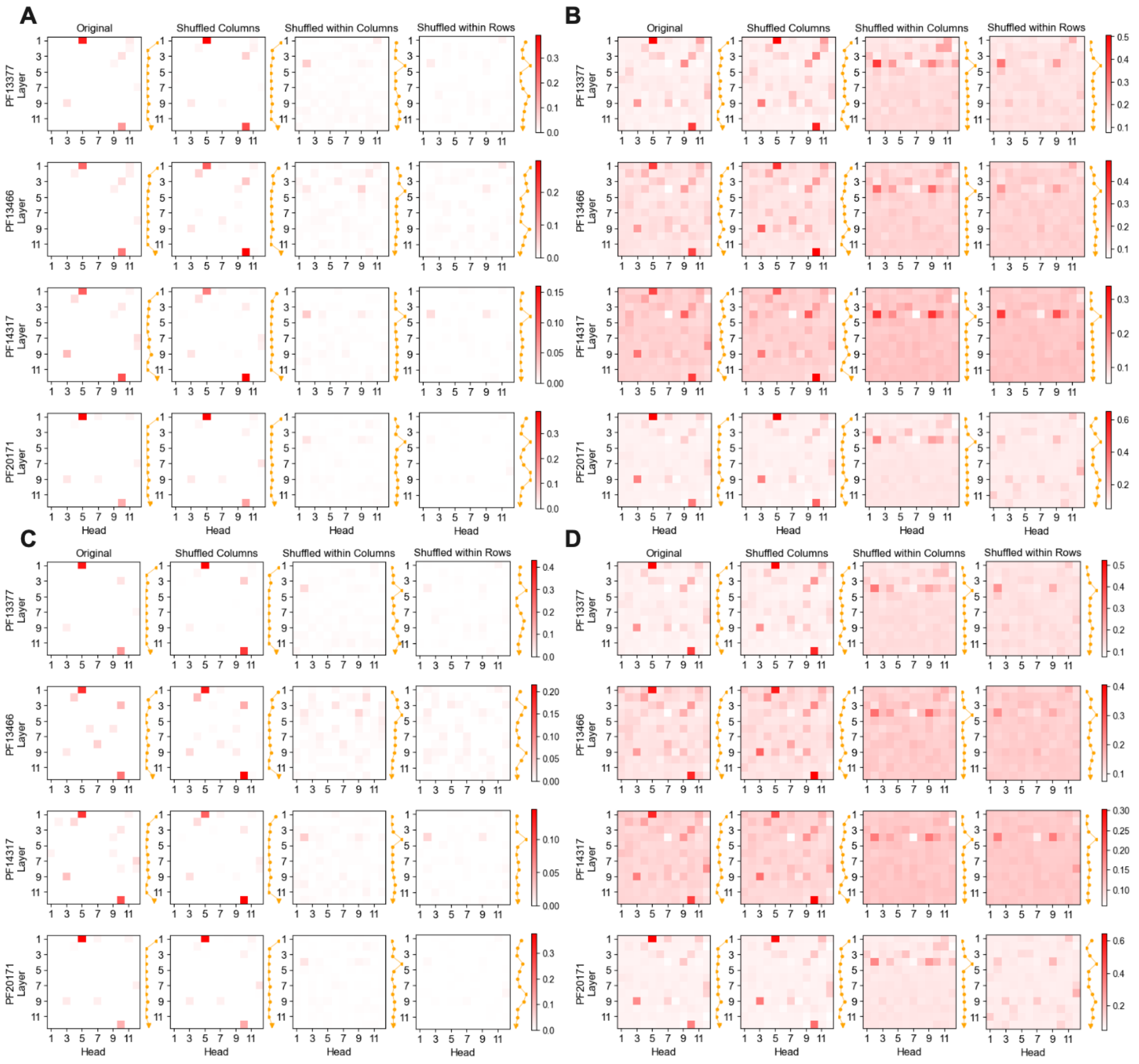


**Figure S9: The heatmap of tree similarity between attention trees and two phylogenetic trees for Pfam protein family *PF13377, PF13466, PF14317, and PF20171*, related to Figure 2.**

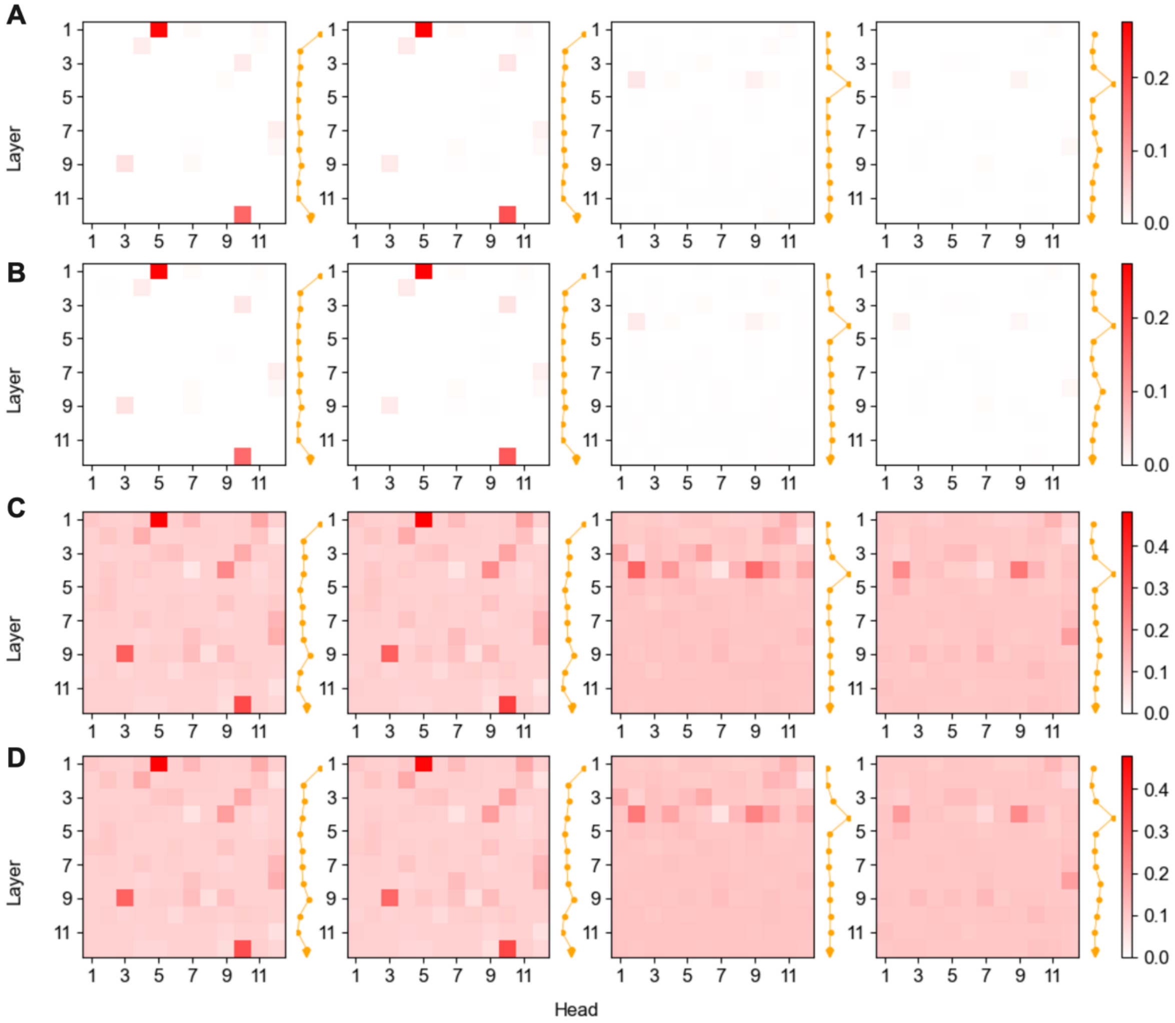


**Figure S10: Consistency in tree similarity between attention trees and two phylogenetic trees evaluated across 20 protein families, related to Figure 2.**
(**A**) Average RF scores between attention trees and the NJ tree.
(**B**) Average RF scores between attention trees and the ML tree.
(**C**) Average CI scores between attention trees and the NJ tree.
(**D**) Average CI scores between attention trees and the ML tree. For details, see Figure 2.

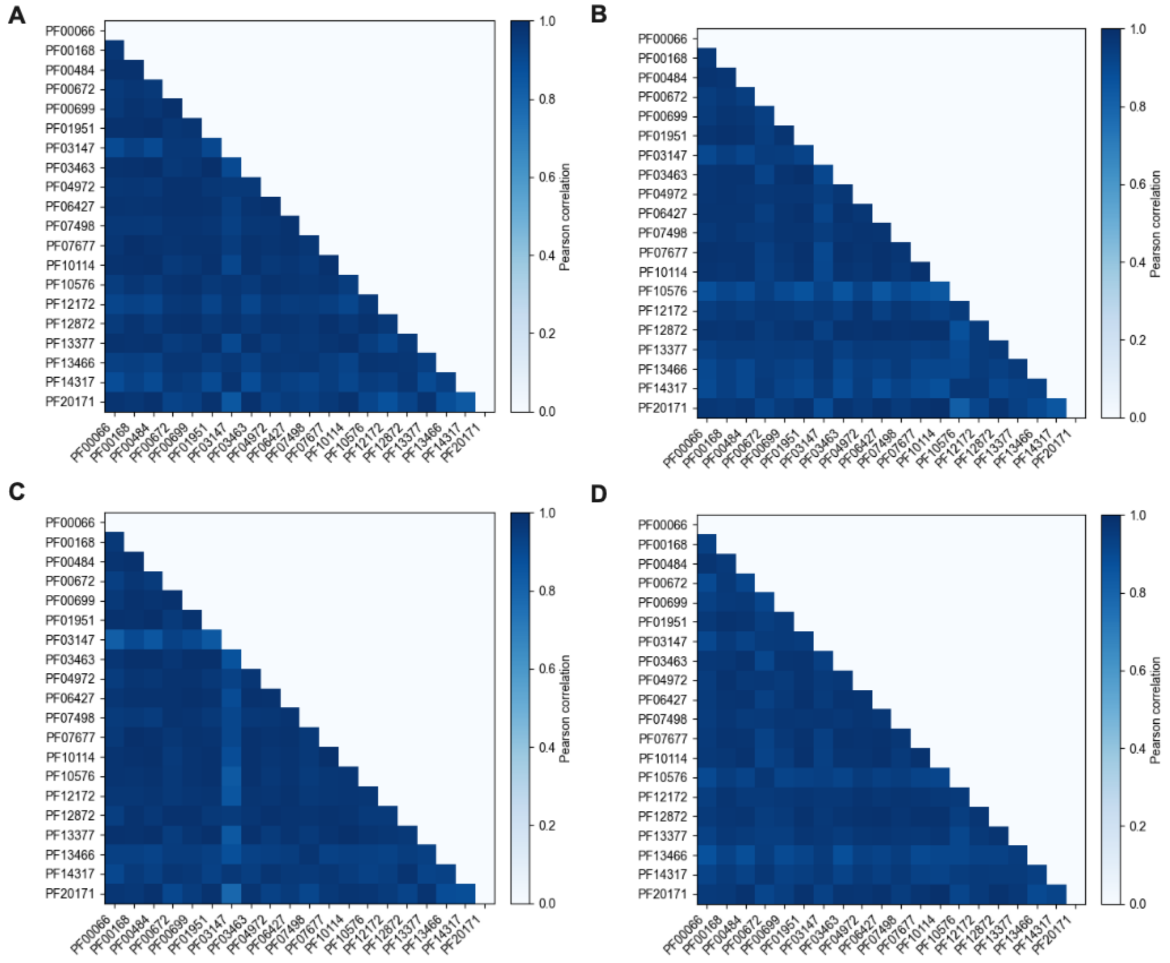


**Figure S11: Pearson correlation analyses of similarity scores across 20 Pfam protein families, related to Figure 2.**
(**A**) Correlation based on RF scores between attention and NJ trees.
(**B**) Correlation based on RF scores between attention and ML trees.
(**C, D**) Equivalent correlation analyses as (A, B), but based on CI scores.

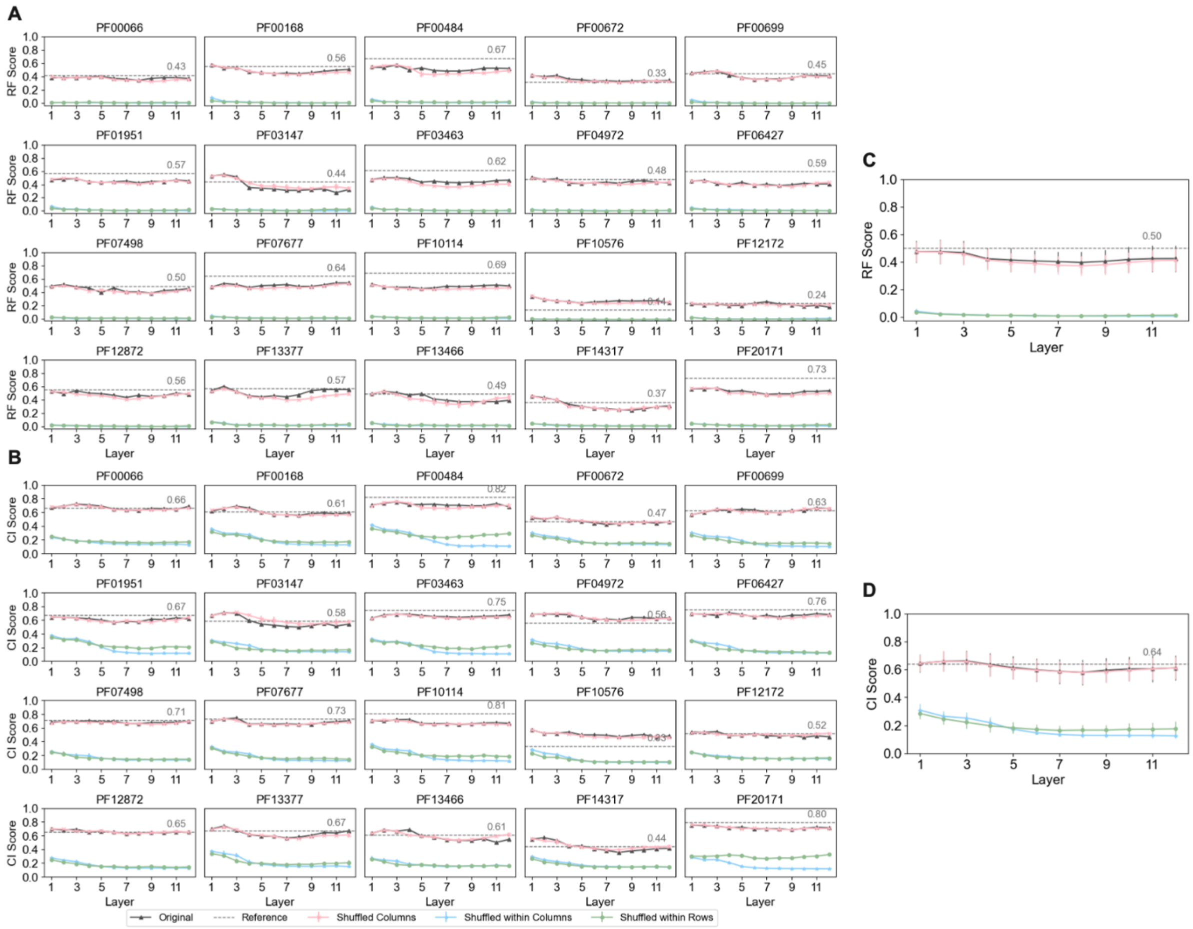


**Figure S12: The similarity scores between embedding trees and *NJ trees*, related to Figure 2.**
(**A, B**) Similarity scores measured by RF and CI scores, respectively. The gray dashed line indicates the reference value between the NJ and ML trees for each protein family. For shuffled MSAs, the y-axis shows the average similarity score from *five shuffling iterations*, with error bars representing standard deviation. (**C, D**) Average RF and CI scores between embedding trees and the NJ tree, respectively. The gray dashed line shows the reference value between NJ and ML trees averaged across *20 protein families*, with error bars indicating standard deviation.

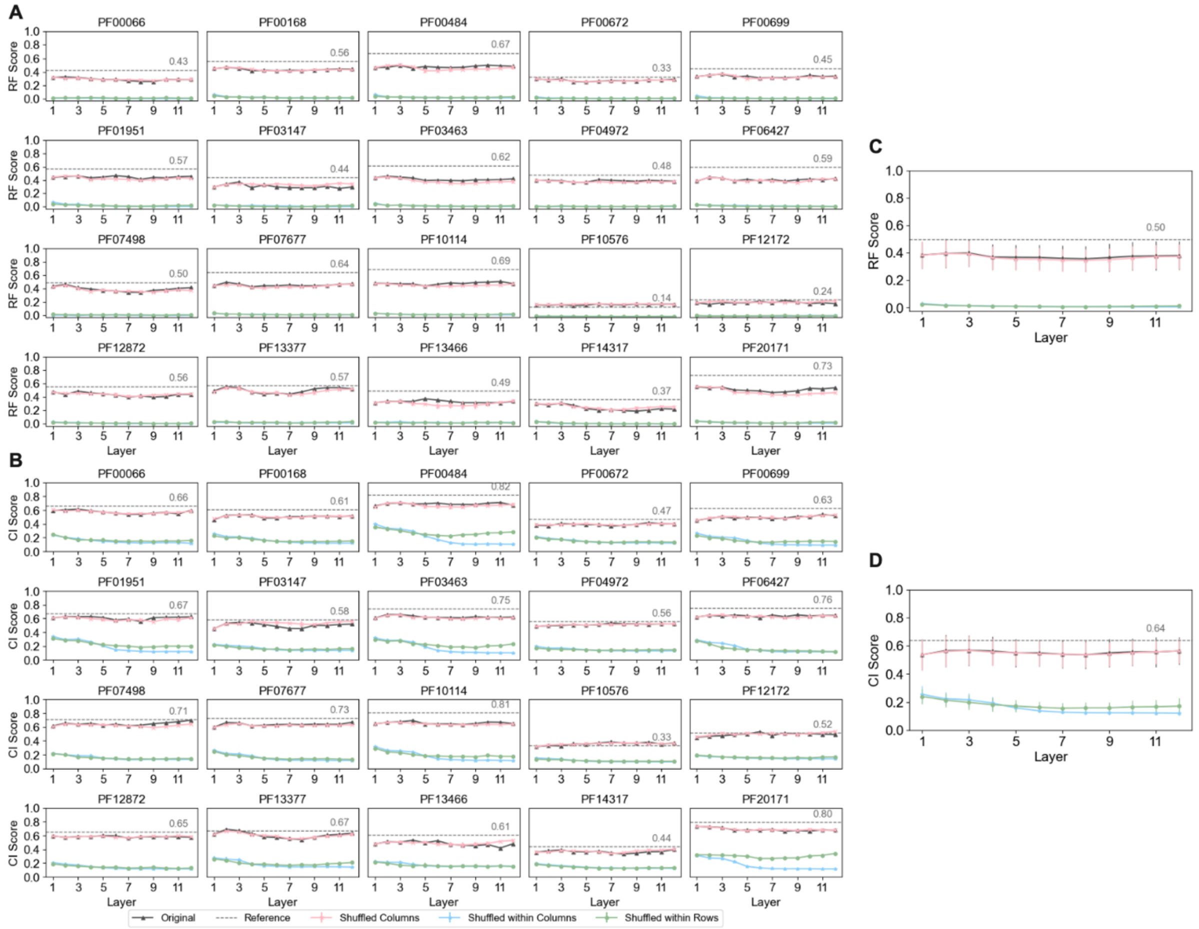


**Figure S13: The similarity scores between embedding trees and *ML trees*, related to Figure 2.**
Same as Figure S12 but for the ML trees.

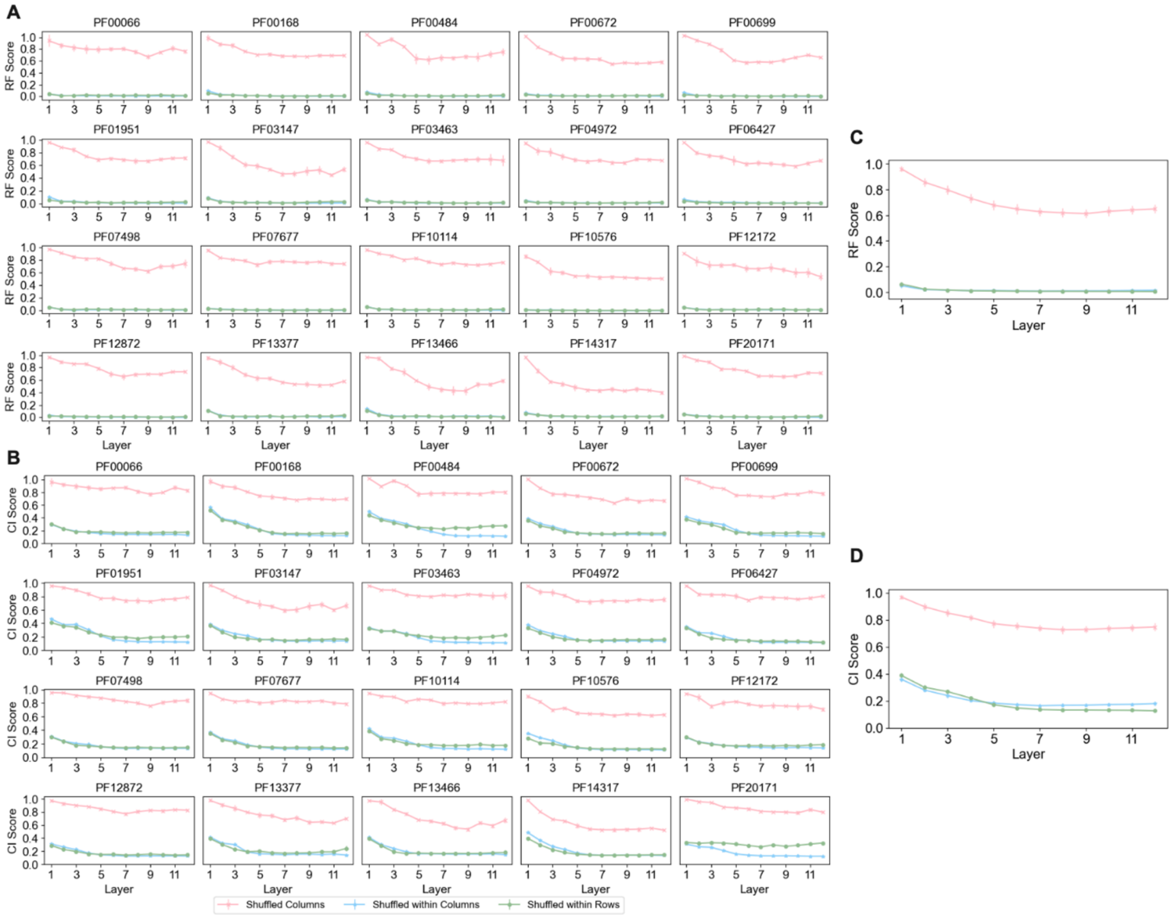


**Figure S14: The tree similarity between embedding tree built from the original MSA and three shuffled MSAs, related to Figure 3.**

(**A**) Layer-by-layer RF score comparisons were performed between embedding trees from original and three types of shuffled MSAs.

(**B**) As in (A), but using CI scores. The y-axis indicates the average similarity score from *five shuffling iterations*, with error bars showing standard deviation.

(**C-D**) Layer-by-layer average (C) RF and (D) CI score comparisons were conducted between embedding trees from original and shuffled MSAs across *20 protein families*, with error bars representing standard deviation.

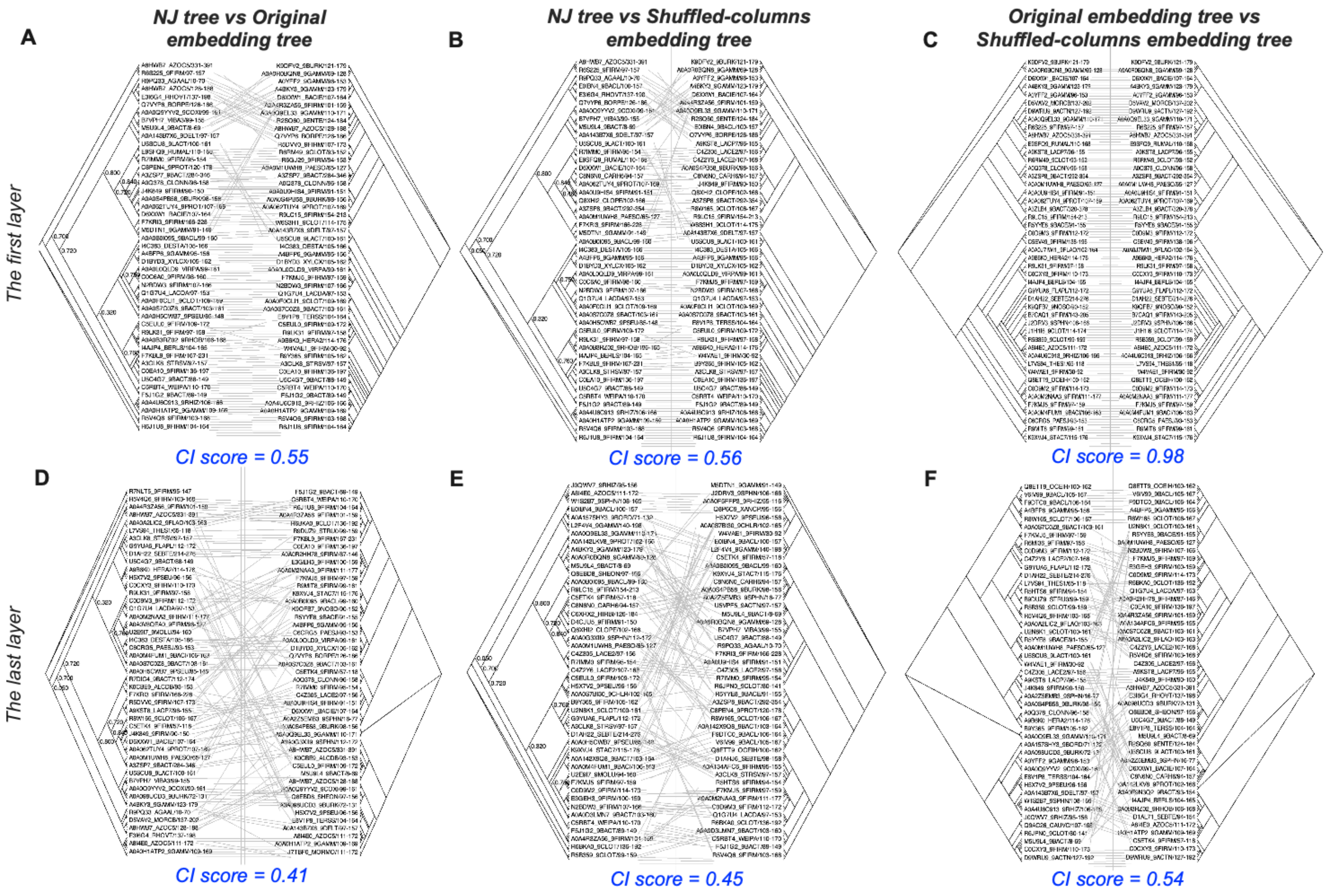


**Figure S15: The tree topology comparison between two trees for Pfam protein family PF14317, related to Figure 3.**
**(A)** Comparison between the NJ tree and the original embedding tree. **(B)** Comparison between the NJ tree and the shuffled-columns embedding tree. **(C)** Comparison between the original embedding tree and the shuffled-columns embedding tree derived from the *first* layer. **(D-F)** Equivalent comparisons for embeddings extracted from the *last* layer. All tree topology comparisons were performed using Dendroscope 3.8.10.[3]

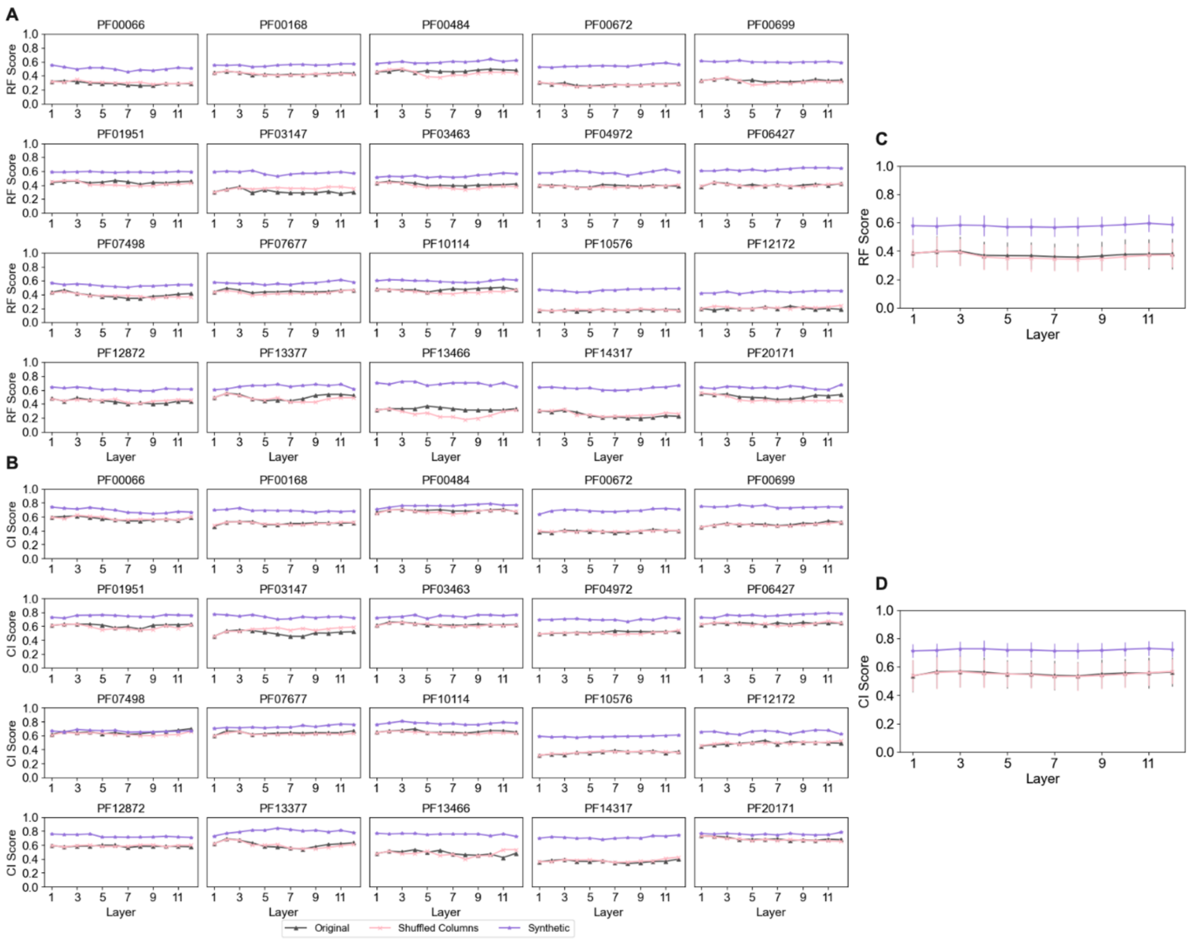


**Figure S16: The similarity between the embedding tree derived from distinct MSAs and their ML trees, related to Figure 3.**

(**A-B**) RF and CI scores were compared between embedding trees generated from original, column-shuffled, and synthetic MSAs, and their corresponding ML trees. ML trees were constructed only from the original Pfam MSAs.

(**C-D**) The average RF and CI scores of embedding trees from different MSAs were compared to those of ML trees across 20 protein families, with error bars representing standard deviation.

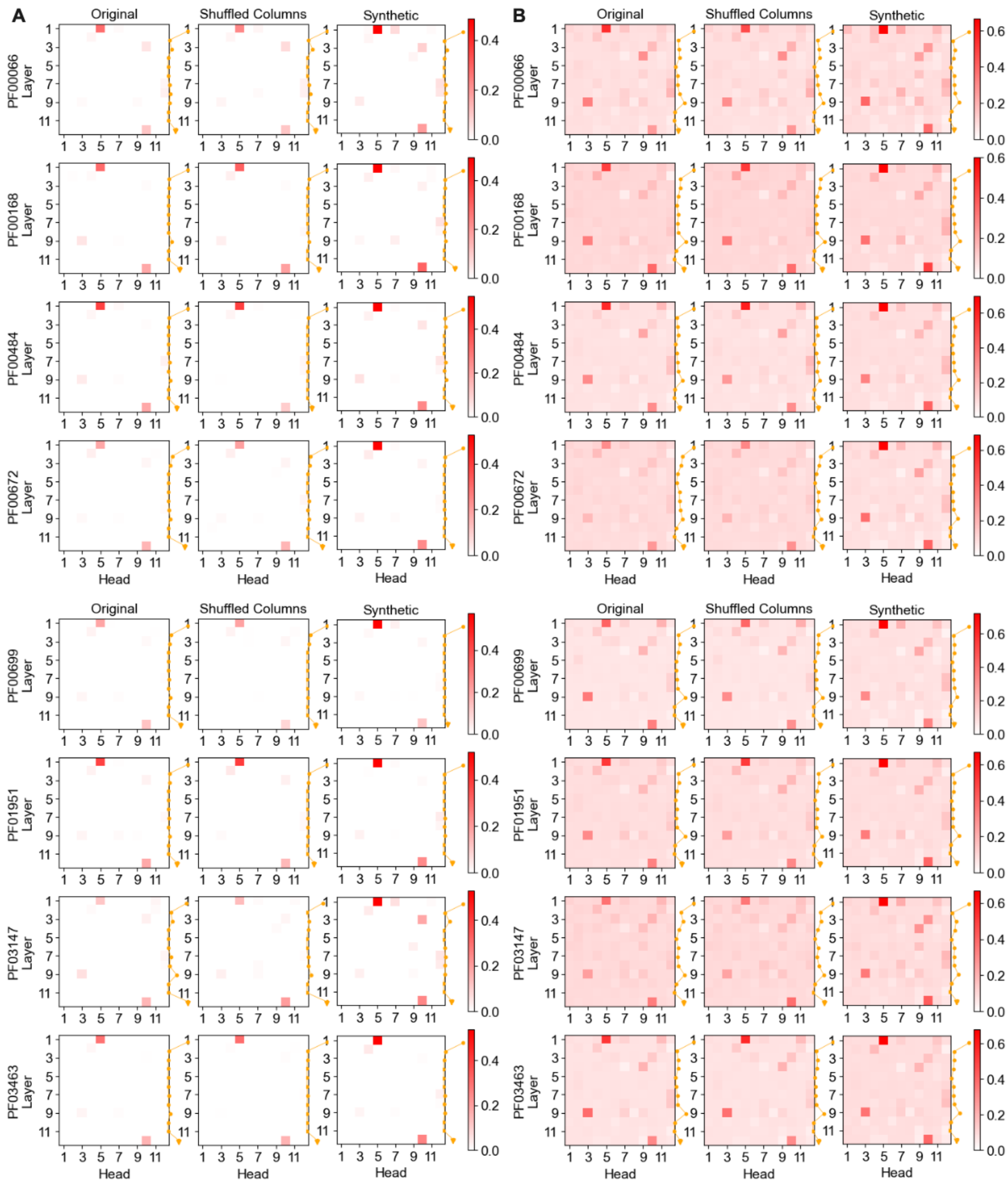


**Figure S17: The tree similarity between attention trees derived from distinct MSAs and their ML trees for *the previous 8* Pfam protein families, related to Figure 3.**
**(A-B)** The RF and CI scores of attention trees derived from original, column-shuffled, and synthetic MSAs were compared with those of their corresponding ML trees for each protein family. For Pfam MSAs, the ML trees were constructed exclusively from the original MSAs.

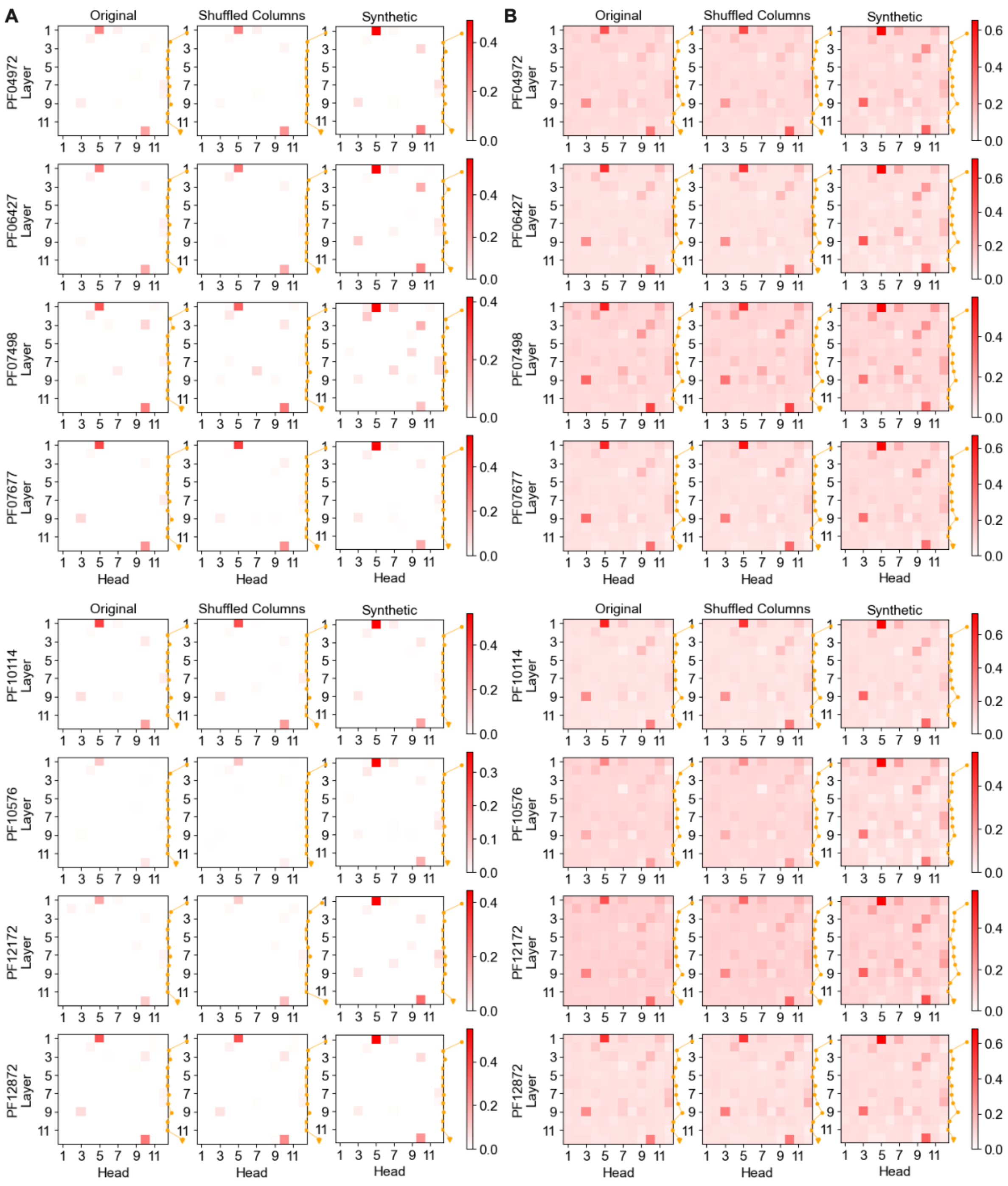


**Figure S18: The tree similarity between attention trees derived from distinct MSAs and their ML trees for *the middle 8* Pfam protein families, related to Figure 3.**

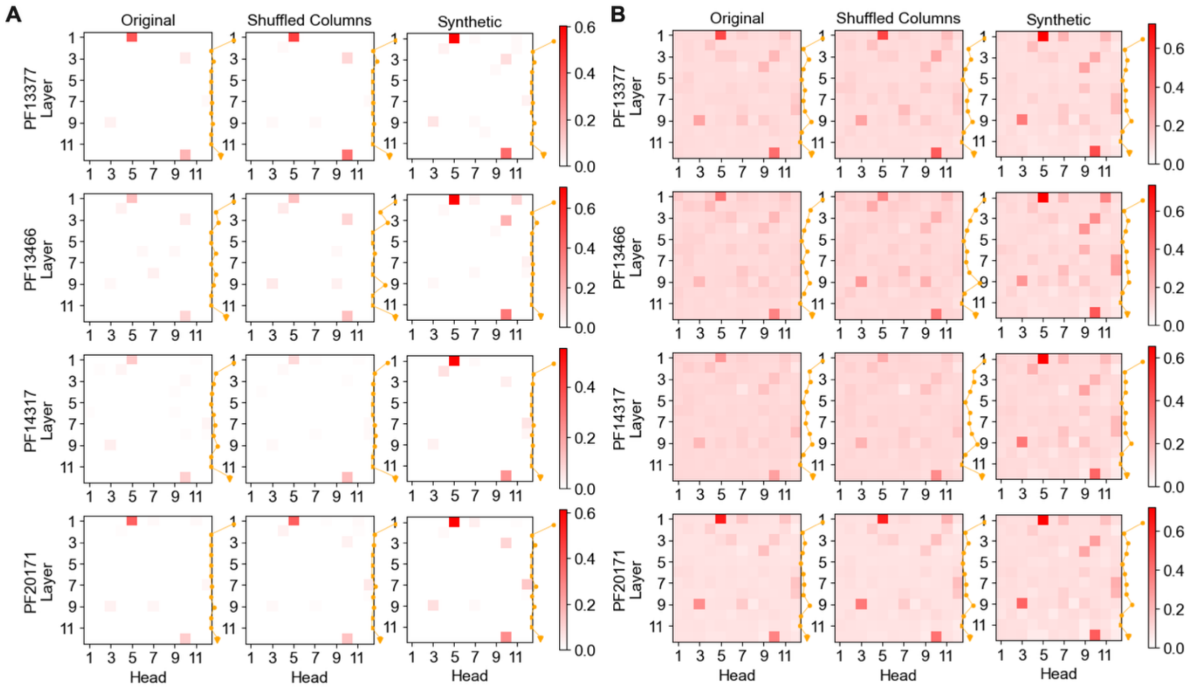


**Figure S19: The tree similarity between attention trees derived from distinct MSAs and their ML trees for *the rest 4* Pfam protein families, related to Figure 3.**

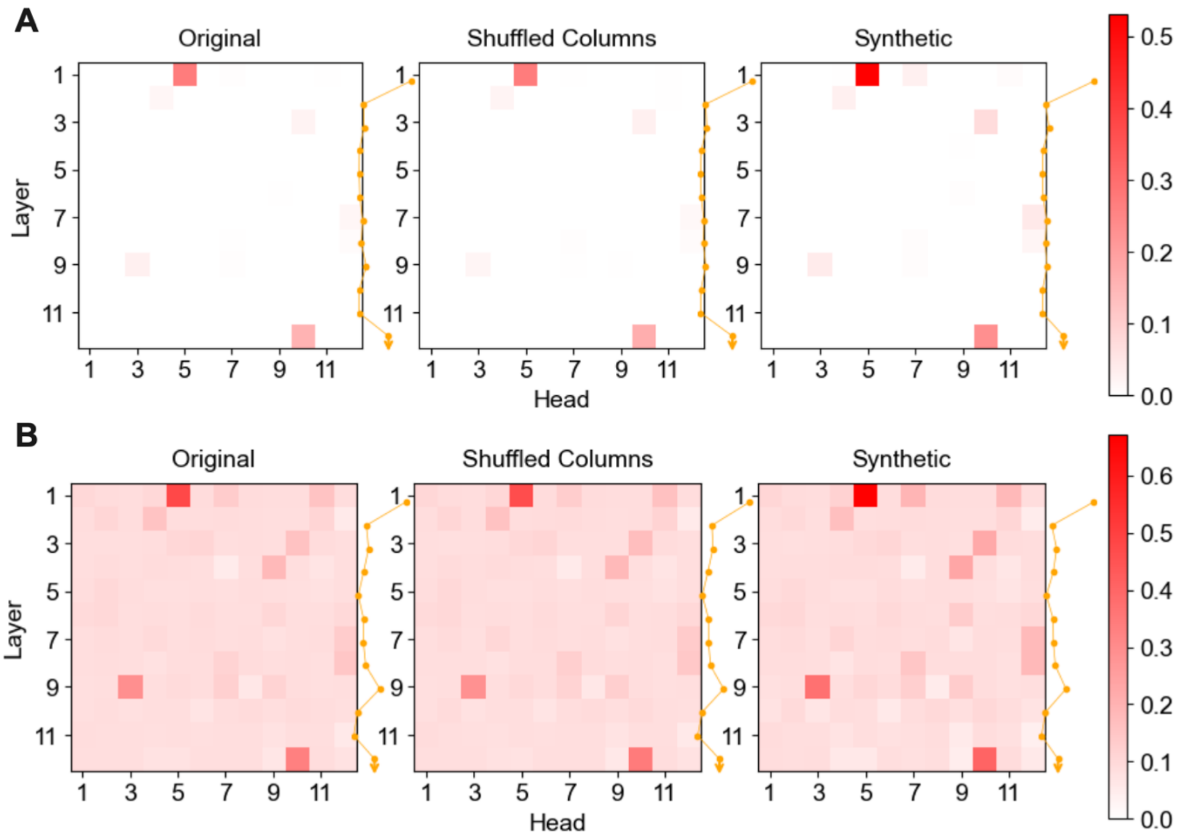


**Figure S20: The average similarity of attention trees derived from original, column-shuffled, and synthetic MSAs were compared with those of their corresponding ML trees across 20 protein families, related to Figure 3.**

(**A-B**) Tree similarity was quantified using RF and CI scores, respectively.

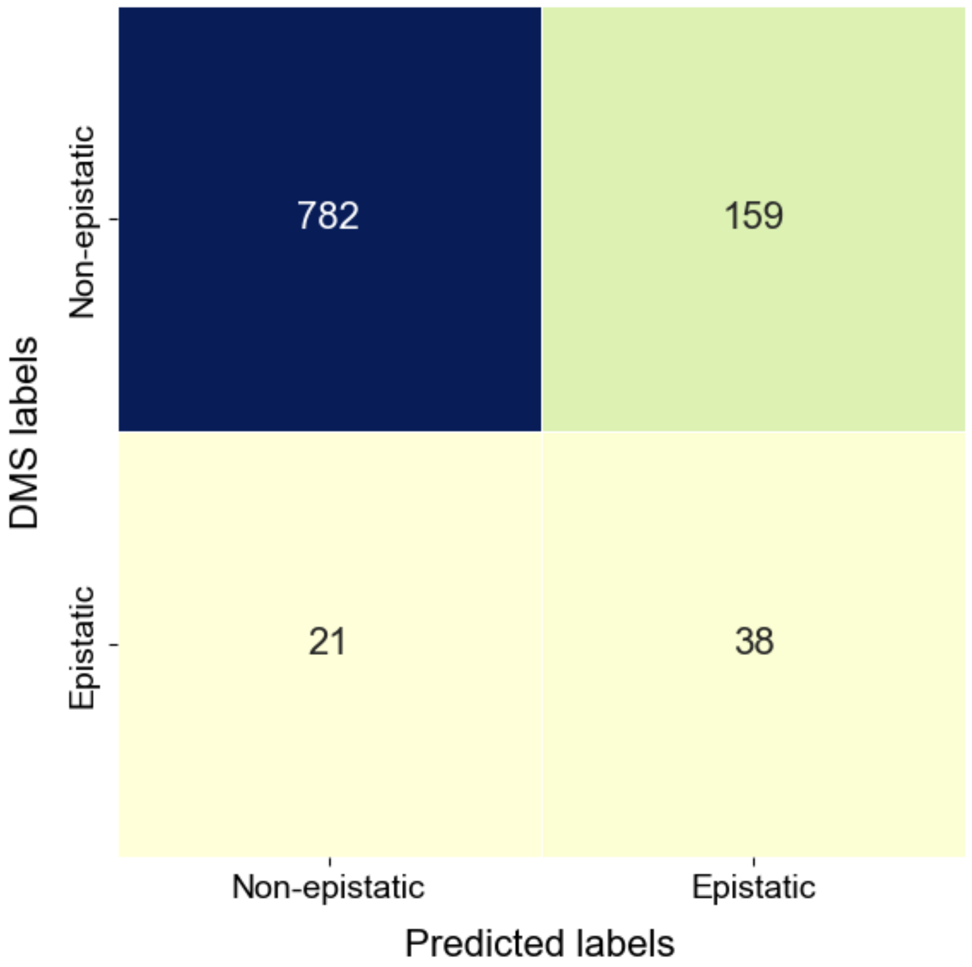


**Figure S21: Confusion matrix comparing predicted and DMS labels for epistatic and non-epistatic variants, related to Figure 3.**
Double mutants with absolute epistasis values exceeding one standard deviation from the mean across 1,000 samples were classified as epistatic, whereas those within one standard deviation were designated as non-epistatic.

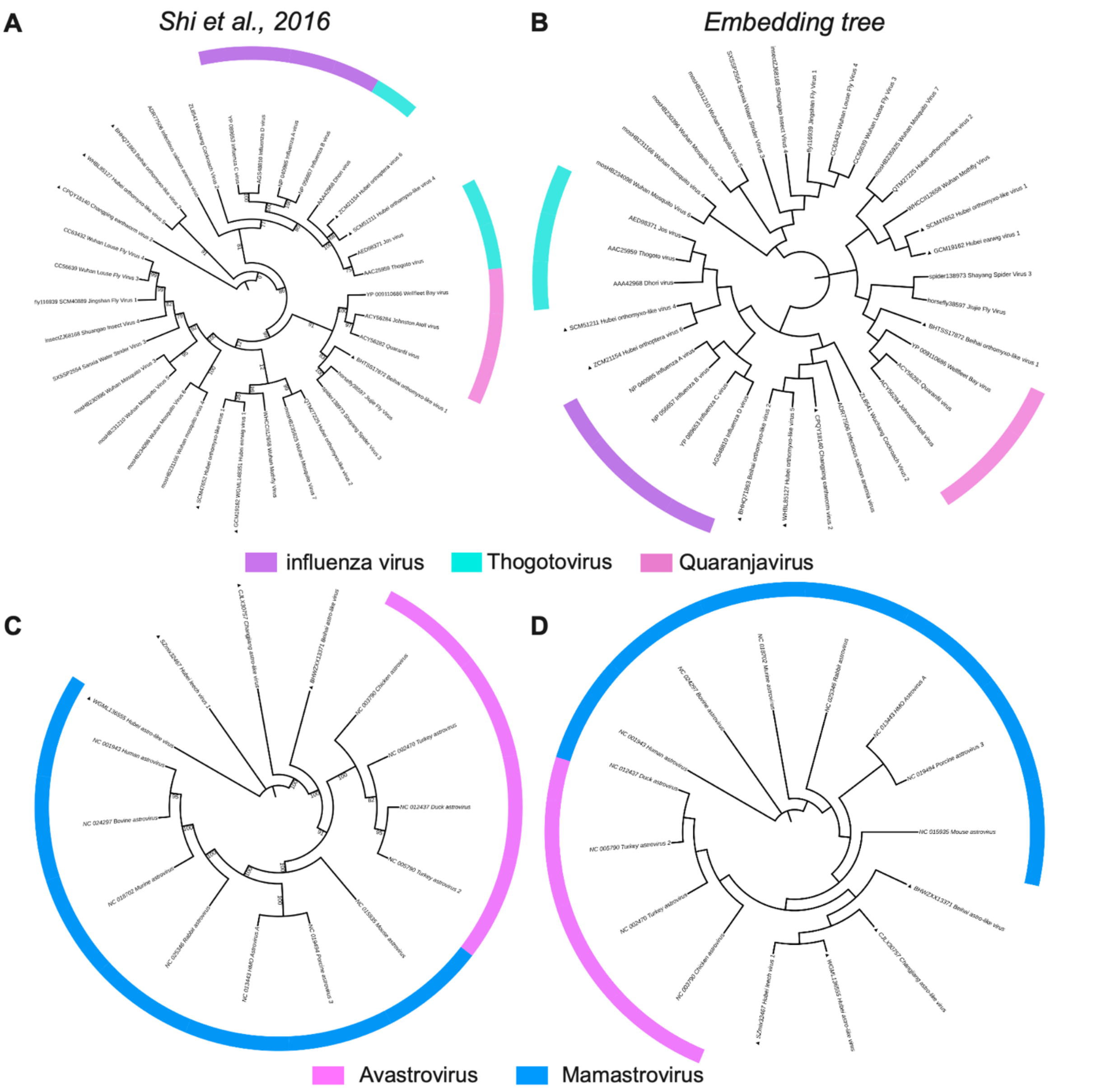


**Figure S22: High consistency between the ML tree and embedding tree for RdRp domain, related to Figure 4.**
(**A-B**) The ML tree and the embedding tree constructed from layer 3 for the "Astro" clade, respectively. (**C-D**) present the same as (A-B), but for the "Orthomyxo" clade. For a fair comparison, the branch lengths were not displayed. Colors represent virus classifications. The trees were visualized using iTOL web tool.[4]

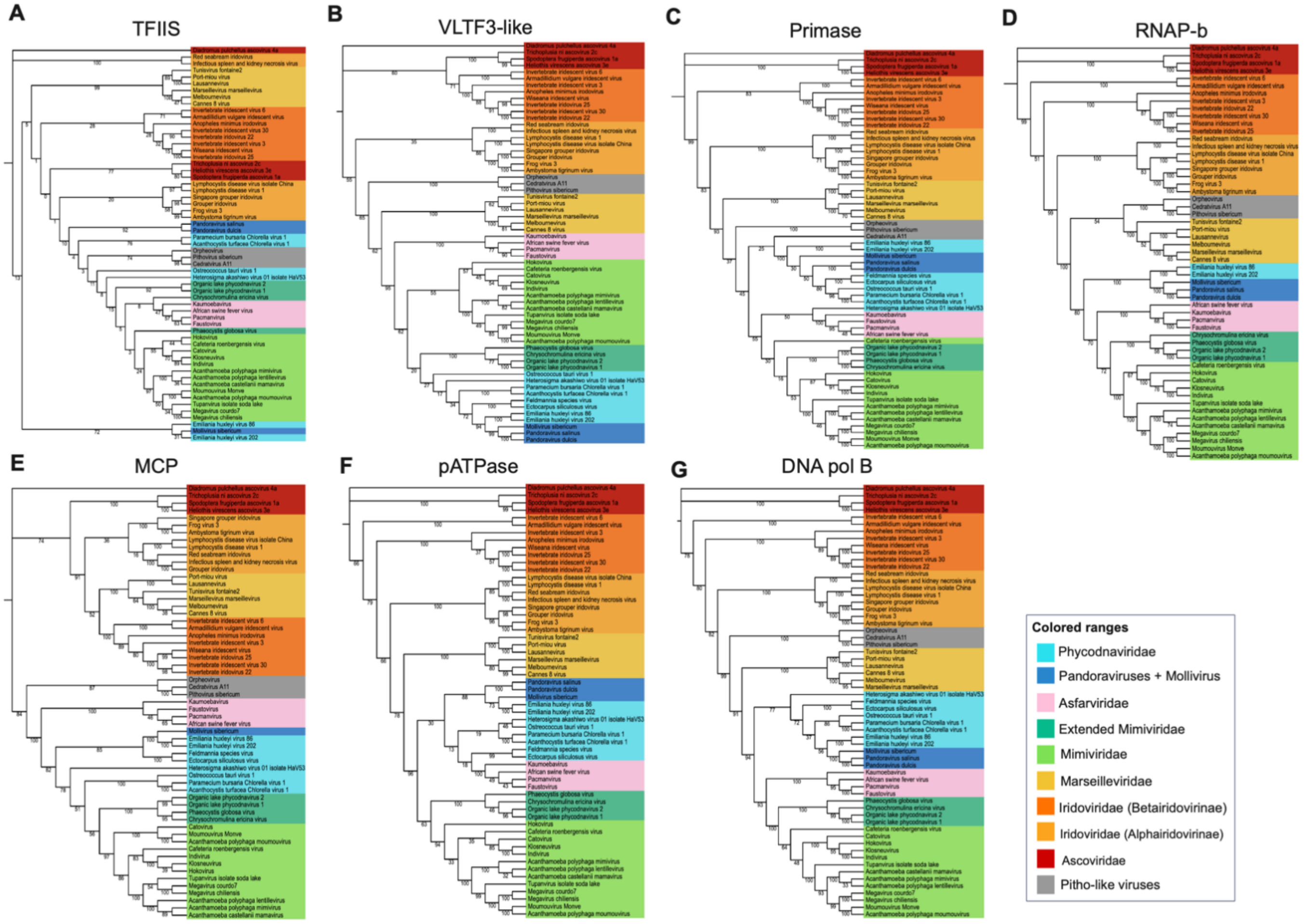


**Figure S23: The ML trees of NCLDV for seven core proteins, related to Figure 5.**
The ML trees for (**A**) TFIIS, (**B**) VLTF3-like, (**C**) Primase, (**D**) RNAP-b, (**E**) MCP, (**F**) pATPase, and (**G**) DNA pol B. For a fair comparison, the branch lengths were not displayed. The trees were visualized using iTOL web tool.[4]

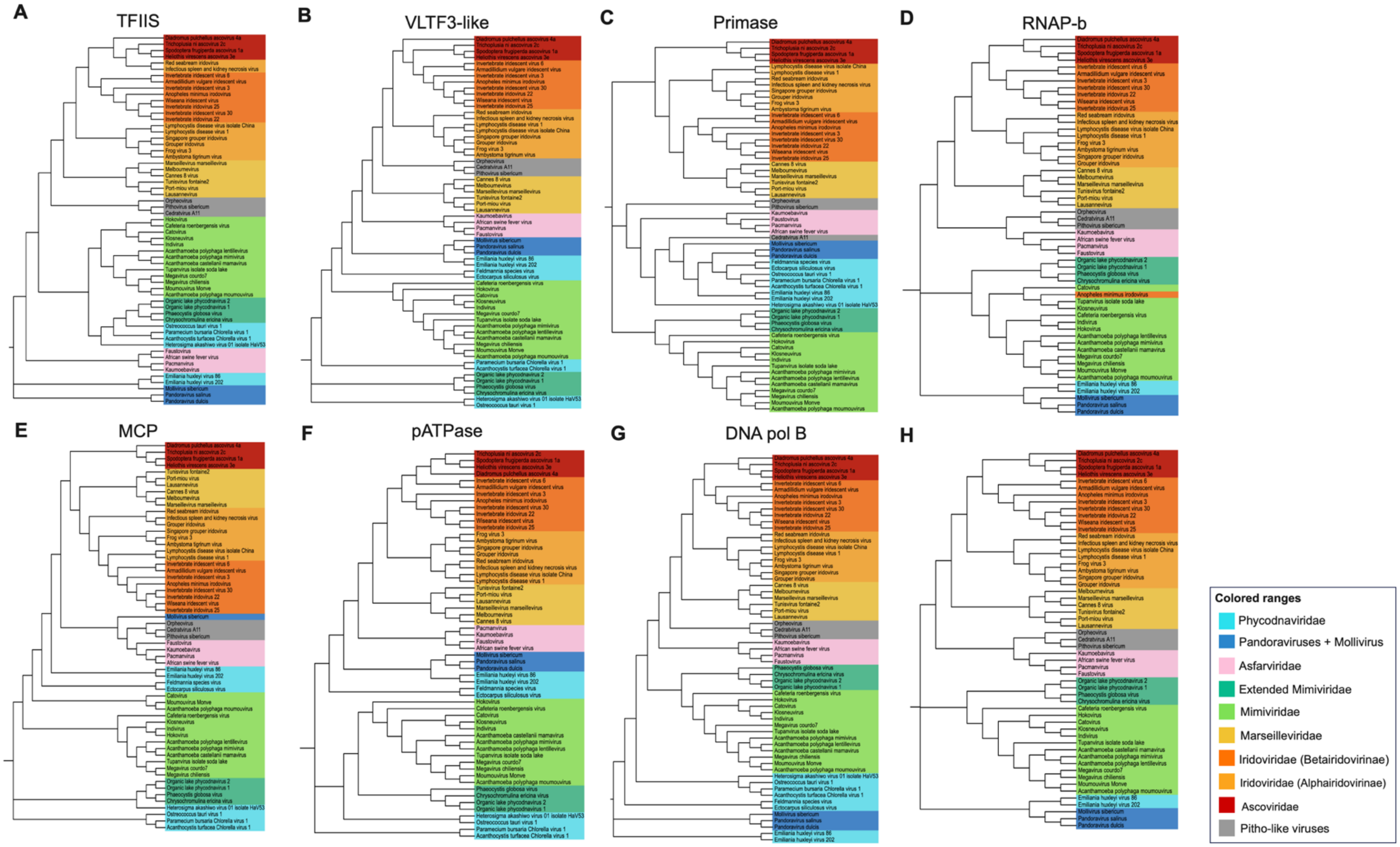


**Figure S24: The embedding trees of NCLDV for seven core proteins, related to Figure 5.**
(**A**) The TFIIS embedding tree constructed from layer 2.
(**B**) The VLTF3-like embedding tree constructed from layer 3.
(**C**) The Primase embedding tree constructed from layer 3.
(**D**) The RNAP-b embedding tree constructed from layer 2.

(**E**) The MCP embedding tree constructed from layer 3.
(**F**) The pATPase embedding tree constructed from layer 2
(**G**) The DNA pol B embedding tree constructed from layer 2.
(**H**) The RNAP-b embedding tree constructed from layer 2 using less gappy MSA. For a fair comparison, the branch lengths were not displayed. The trees were visualized using iTOL web tool.[4]

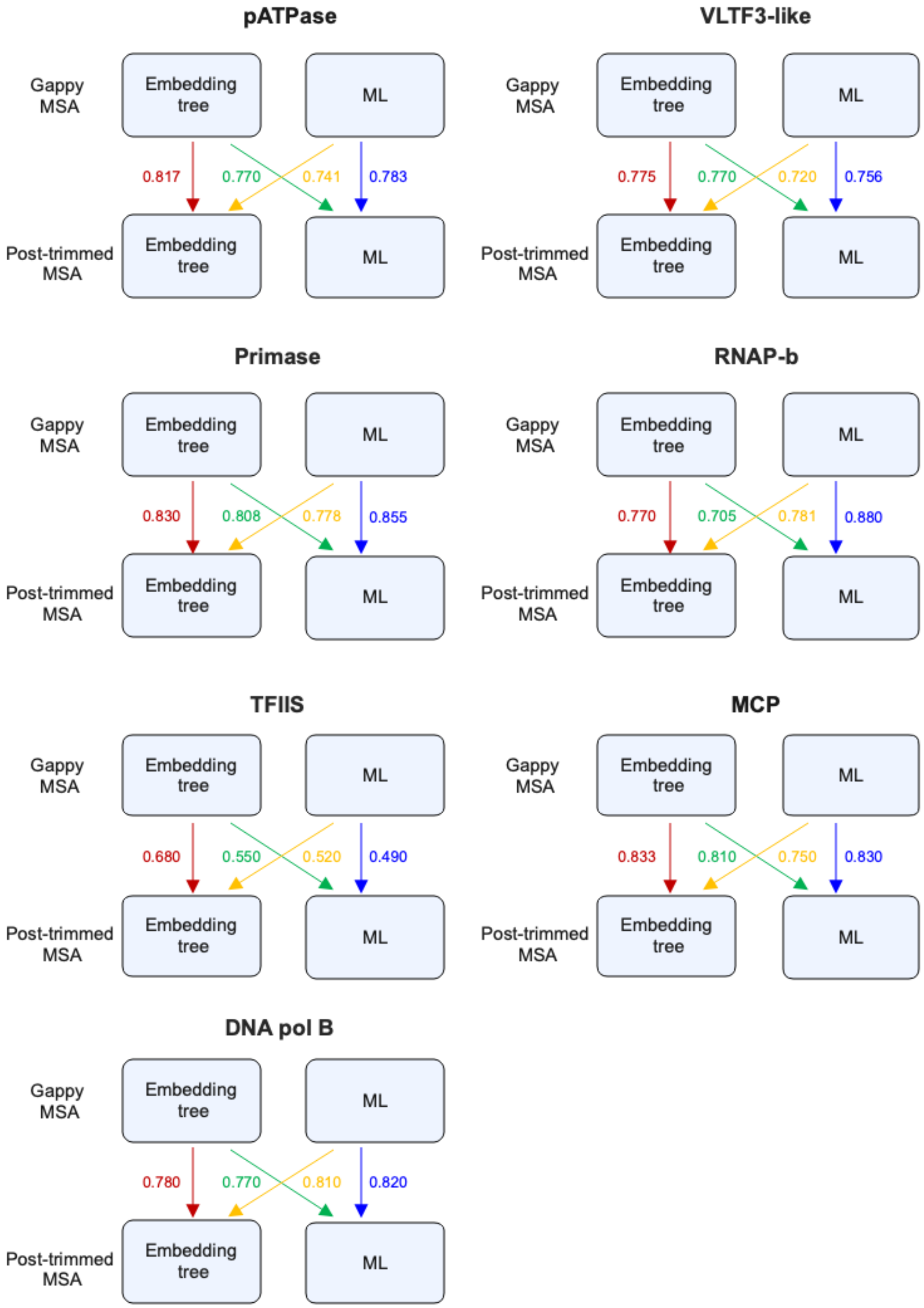


**Figure S25: The ability of MSA Transformer and ML method to recover phylogenetic relationships from gappy MSA, related to Figure 5.**
Four different comparisons were indicated using distinct colors. The value next to the arrow for each comparison is the average CI score calculated over 50 replicates.

**Supplemental tables**

| | Pfam ID | Protein family | $M$ | $L$ | $\mu_L$ | $\delta$ (NJ) | $\delta$ (ML) |
|---|---|---|---|---|---|---|---|
| 1 | PF00066 | LNR domain | 104 | 40 | 35.88 | 2.03 | 1.86 |
| 2 | PF00168 | C2 domain | 258 | 253 | 107.60 | 1.95 | 2.67 |
| 3 | PF00484 | Carbonic anhydrase | 252 | 285 | 155.87 | 2.44 | 2.47 |
| 4 | PF00672 | HAMP domain | 212 | 80 | 53.84 | 2.04 | 2.17 |
| 5 | PF00699 | Urease beta subunit | 294 | 145 | 98.31 | 1.10 | 1.47 |
| 6 | PF01951 | Archease protein domain | 192 | 285 | 137.64 | 1.75 | 1.68 |
| 7 | PF03147 | Ferredoxin-fold anticodon binding domain | 96 | 134 | 94.14 | 1.35 | 1.75 |
| 8 | PF03463 | eRF1 domain | 242 | 288 | 129.00 | 2.24 | 2.59 |
| 9 | PF04972 | BON domain | 138 | 90 | 68.33 | 1.99 | 2.12 |
| 10 | PF06427 | UDP-glucose:Glycoprotein Glucosyltransferase | 155 | 136 | 109.23 | 1.37 | 1.53 |
| 11 | PF07498 | Rho termination factor, N-terminal domain | 104 | 50 | 42.84 | 1.08 | 1.11 |
| 12 | PF07677 | A-macroglobulin receptor binding domain | 194 | 130 | 93.03 | 1.95 | 2.02 |
| 13 | PF10114 | Sensory domain found in PocR | 195 | 242 | 162.35 | 2.07 | 2.59 |
| 14 | PF10576 | Iron-sulfur binding domain of endonuclease III | 392 | 23 | 17.02 | 1.95 | 1.53 |
| 15 | PF12172 | Rubredoxin-like zinc ribbon domain | 93 | 43 | 36.83 | 2.10 | 2.08 |
| 16 | PF12872 | LOTUS domain | 130 | 105 | 71.45 | 2.91 | 3.65 |
| 17 | PF13377 | Periplasmic binding protein-like domain | 66 | 264 | 164.34 | 2.14 | 2.32 |
| 18 | PF13466 | STAS domain | 54 | 101 | 79.89 | 1.90 | 2.23 |
| 19 | PF14317 | YcxB-like domain | 141 | 101 | 61.31 | 2.20 | 2.80 |
| 20 | PF20171 | Glucose-6-phosphate dehydrogenase subunit C-terminal domain | 130 | 275 | 152.12 | 2.05 | 2.50 |

**Table S1: Pfam MSAs and their associated phylogenetic trees are used in this work, related to Figure 2.**

| Types of MSA | Retained from original | Disrupted from original | Biology |
|---|---|---|---|
| Original | N/A | N/A | Homologous positions, potential epistasis |
| Shuffled-columns | Row index and composition | Column re-indexed | Homologous positions, potential epistasis |
| Shuffled-within-columns | Column index and composition | Row re-indexed for each column | Scrambled MSA |
| Shuffled-within-rows | MSA and row composition | Column re-indexed for each row | Scrambled MSA |
| Synthetic MSA | N/A | N/A | Fictional homology across sequence, no epistasis |

**Table S2: Five scenarios for data analysis of MSA, designed to probe dependency of column and row-format, and sensitivity to potential epistasis, related to Figure 3.**

| | Synthetic ID | *Extant* | *Models* | *Rates* | *Dist* |
|---|---|---|---|---|---|
| 1 | SPF00066 | 104 | LG | 0.01 | 1.30 |
| 2 | SPF00168 | 258 | LG | 0.05 | 2.10 |
| 3 | SPF00484 | 252 | LG | 0.03 | 1.40 |
| 4 | SPF00672 | 212 | LG | 0.02 | 1.55 |
| 5 | SPF00699 | 294 | LG | 0.02 | 1.00 |
| 6 | SPF01951 | 192 | LG | 0.04 | 1.10 |
| 7 | SPF03147 | 96 | LG | 0.01 | 1.55 |
| 8 | SPF03463 | 242 | LG | 0.04 | 1.60 |
| 9 | SPF04972 | 138 | LG | 0.01 | 1.60 |
| 10 | SPF06427 | 155 | LG | 0.009 | 1.15 |
| 11 | SPF07498 | 104 | LG | 0.01 | 0.80 |
| 12 | SPF07677 | 194 | LG | 0.01 | 1.60 |
| 13 | SPF10114 | 195 | LG | 0.01 | 1.65 |
| 14 | SPF10576 | 392 | LG | 0.01 | 1.45 |
| 15 | SPF12172 | 93 | LG | 0.01 | 1.85 |
| 16 | SPF12872 | 130 | LG | 0.02 | 2.65 |
| 17 | SPF13377 | 66 | LG | 0.035 | 1.635 |
| 18 | SPF13466 | 54 | LG | 0.02 | 1.55 |
| 19 | SPF14317 | 141 | LG | 0.024 | 2.10 |
| 20 | SPF20171 | 130 | LG | 0.02 | 1.85 |

**Table S3: Parameters employed by TrAVIS[5] to generate Synthetic MSAs, related to Figure 3.**

| | Pfam ID | $M$ | The original MSA | | Synthetic MSA | |
|---|---|---|---|---|---|---|
| | | | $L$ | $\delta$ | $L$ | $\delta$ |
| 1 | PF00066 | 104 | 40 | 1.86 | 41 | 1.84 |
| 2 | PF00168 | 258 | 253 | 2.67 | 252 | 2.67 |
| 3 | PF00484 | 252 | 285 | 2.47 | 291 | 2.46 |
| 4 | PF00672 | 212 | 80 | 2.17 | 76 | 2.15 |
| 5 | PF00699 | 294 | 145 | 1.47 | 148 | 1.50 |
| 6 | PF01951 | 192 | 285 | 1.68 | 281 | 1.74 |
| 7 | PF03147 | 96 | 134 | 1.75 | 138 | 1.74 |
| 8 | PF03463 | 242 | 288 | 2.59 | 281 | 2.51 |
| 9 | PF04972 | 138 | 90 | 2.12 | 96 | 2.09 |
| 10 | PF06427 | 155 | 136 | 1.53 | 134 | 1.46 |
| 11 | PF07498 | 104 | 50 | 1.11 | 52 | 1.10 |
| 12 | PF07677 | 194 | 130 | 2.02 | 139 | 1.97 |
| 13 | PF10114 | 195 | 242 | 2.59 | 238 | 2.54 |
| 14 | PF10576 | 392 | 23 | 1.53 | 21 | 1.56 |
| 15 | PF12172 | 93 | 43 | 2.08 | 47 | 2.05 |
| 16 | PF12872 | 130 | 105 | 3.65 | 113 | 3.59 |
| 17 | PF13377 | 66 | 264 | 2.32 | 264 | 2.29 |
| 18 | PF13466 | 54 | 101 | 2.23 | 103 | 2.21 |
| 19 | PF14317 | 141 | 101 | 2.80 | 101 | 2.86 |
| 20 | PF20171 | 130 | 275 | 2.50 | 278 | 2.48 |

**Table S4: Comparison of the original MSA, synthetic MSA and their ML tree, related to Figure 3.**

| | Name | $M$ | $L$ | $\delta$ |
|---|---|---|---|---|
| 1 | Reo | 86 | 414 | 410.05 |
| 2 | Nido | 36 | 421 | 412.47 |
| 3 | Orthomyxo | 34 | 307 | 305.09 |
| 4 | Astro | 15 | 321 | 317.33 |

**Table S5: Four clades for RdRps used in this work, related to Figure 4.**

| Layer | Astro | | Orthomyxo | | Nido | | Reo | |
|---|---|---|---|---|---|---|---|---|
| | RF | CI | RF | CI | RF | CI | RF | CI |
| 1 | 0.50 | 0.77 | 0.81 | 0.84 | 0.69 | 0.78 | 0.68 | 0.82 |
| 2 | 1.00 | 1.00 | 0.77 | 0.80 | 0.72 | 0.80 | 0.71 | 0.82 |
| 3 | 1.00 | 1.00 | 0.84 | 0.83 | 0.72 | 0.78 | 0.77 | 0.83 |
| 4 | 0.92 | 0.91 | 0.84 | 0.83 | 0.62 | 0.71 | 0.68 | 0.78 |
| 5 | 0.92 | 0.91 | 0.84 | 0.83 | 0.66 | 0.74 | 0.67 | 0.78 |
| 6 | 0.92 | 0.91 | 0.84 | 0.83 | 0.56 | 0.65 | 0.67 | 0.79 |
| 7 | 0.92 | 0.91 | 0.77 | 0.79 | 0.56 | 0.65 | 0.60 | 0.75 |
| 8 | 0.92 | 0.91 | 0.74 | 0.78 | 0.56 | 0.65 | 0.63 | 0.75 |
| 10 | 0.92 | 0.91 | 0.77 | 0.80 | 0.56 | 0.65 | 0.63 | 0.74 |
| 11 | 0.83 | 0.89 | 0.81 | 0.85 | 0.66 | 0.72 | 0.64 | 0.75 |
| 12 | 0.92 | 0.95 | 0.81 | 0.82 | 0.69 | 0.74 | 0.66 | 0.77 |

**Figure S6: The similarity scores between embedding tree and ML tree for NCLDVs, related to Figure 4.**

The specific layer chosen to build the embedding tree for each protein was marked by yellow.

| | Name | $M$ | $L$ | | $\delta$ |
|---|---|---|---|---|---|
| | | | Pre-trimmed | Post-trimmed | |
| 1 | DNA pol B | 61 | 4805 | 841 | 827.11 |
| 2 | TFIIS | 59 | 527 | 75 | 72.36 |
| 3 | MCP | 59 | 1447 | 397 | 384.03 |
| 4 | pATPase | 58 | 831 | 234 | 231.09 |
| 5 | Primase | 61 | 2257 | 744 | 697.48 |
| 6 | VLTF3-like | 61 | 994 | 283 | 265.34 |
| 7 | RNAP-b | 55 | 3795 | 1022 | 983.90 |

**Figure S7: Seven core proteins for NCLDVs used in this work, related to Figure 5.**

| Layer | MCP | | DNA pol B | | pATPase | | Primase | |
|---|---|---|---|---|---|---|---|---|
| | RF | CI | RF | CI | RF | CI | RF | CI |
| 1 | 0.66 | 0.77 | 0.69 | 0.80 | 0.80 | 0.81 | 0.69 | 0.82 |
| 2 | 0.73 | 0.78 | 0.83 | 0.87 | 0.84 | 0.83 | 0.83 | 0.85 |
| 3 | 0.70 | 0.76 | 0.81 | 0.83 | 0.85 | 0.90 | 0.83 | 0.83 |
| 4 | 0.70 | 0.75 | 0.62 | 0.66 | 0.80 | 0.83 | 0.76 | 0.70 |
| 5 | 0.68 | 0.73 | 0.69 | 0.74 | 0.82 | 0.88 | 0.76 | 0.70 |
| 6 | 0.64 | 0.71 | 0.60 | 0.62 | 0.75 | 0.82 | 0.76 | 0.71 |
| 7 | 0.61 | 0.63 | 0.59 | 0.59 | 0.73 | 0.81 | 0.69 | 0.65 |
| 8 | 0.59 | 0.60 | 0.55 | 0.57 | 0.69 | 0.76 | 0.71 | 0.65 |
| 9 | 0.61 | 0.69 | 0.57 | 0.60 | 0.71 | 0.75 | 0.64 | 0.63 |
| 10 | 0.54 | 0.52 | 0.53 | 0.55 | 0.69 | 0.74 | 0.64 | 0.63 |
| 11 | 0.54 | 0.51 | 0.59 | 0.62 | 0.55 | 0.64 | 0.69 | 0.66 |
| 12 | 0.59 | 0.57 | 0.60 | 0.64 | 0.73 | 0.82 | 0.71 | 0.65 |

| Layer | RNAP-b | | TFIIS | | VLTF3-like | |
|---|---|---|---|---|---|---|
| | RF | CI | RF | CI | RF | CI |
| 1 | 0.62 | 0.75 | 0.57 | 0.58 | 0.62 | 0.76 |
| 2 | 0.63 | 0.78 | 0.63 | 0.67 | 0.69 | 0.77 |
| 3 | 0.63 | 0.75 | 0.63 | 0.66 | 0.79 | 0.81 |
| 4 | 0.60 | 0.63 | 0.55 | 0.59 | 0.78 | 0.80 |
| 5 | 0.62 | 0.63 | 0.57 | 0.59 | 0.72 | 0.75 |
| 6 | 0.63 | 0.65 | 0.55 | 0.56 | 0.72 | 0.73 |
| 7 | 0.62 | 0.64 | 0.55 | 0.53 | 0.66 | 0.67 |
| 8 | 0.63 | 0.65 | 0.54 | 0.49 | 0.59 | 0.59 |
| 9 | 0.60 | 0.58 | 0.54 | 0.50 | 0.52 | 0.54 |
| 10 | 0.60 | 0.60 | 0.52 | 0.50 | 0.52 | 0.55 |
| 11 | 0.62 | 0.63 | 0.52 | 0.49 | 0.59 | 0.62 |
| 12 | 0.63 | 0.67 | 0.55 | 0.56 | 0.64 | 0.68 |

**Figure S8: The similarity scores between the embedding tree and ML tree of seven core proteins for NCLDVs, related to Figure 5.**

The specific layer chosen to build the embedding tree for each protein was marked by yellow.
